## Supplementary Information containing Sup. Fig.1-14 for "SDR enzymes oxidize specific lipidic alkynylcarbinols into cytotoxic protein-reactive species"

This file includes:

### Supplementary figures 1 to 13.

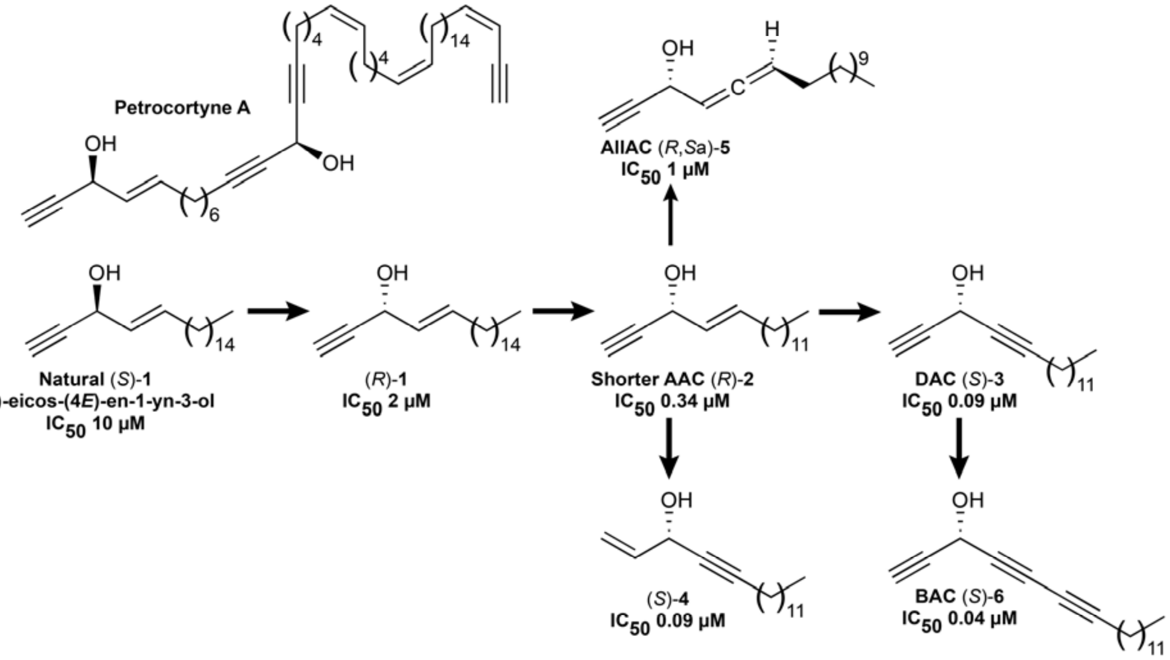

**Supplementary Figure 1: Representative structures of natural and bioinspired synthetic alkynylcarbinol-containing cytotoxic molecules.** Previously reported $IC_{50}$ values, evaluated in HCT116 cells, are indicated when available.

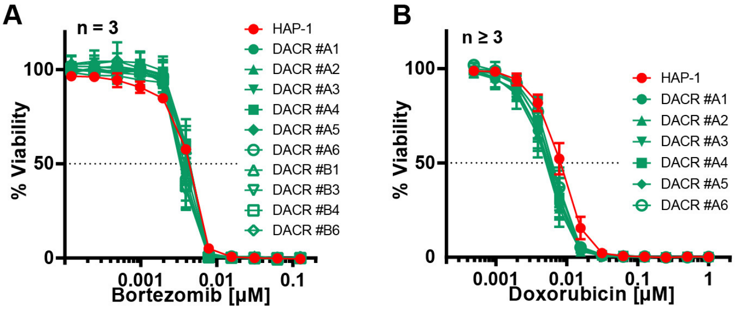

C

| Clone | Gene | Mutation |
|---|---|---|
| A1 | ACOX3 | His576Asp |
| | ACSF2 | Lys599Asn |
| | ATG9A | Pro660Ser |
| | CDC23 | Val177Ala |
| | GPC2 | Arg147Trp |
| | **HSD17B11** | Val16Asp |
| | **KCTD5** | Glu196* |
| | MYDGF | Gly22Arg |
| | PCNX3 | Pro1117Leu |
| | WWTR1 | Asn213Tyr |
| A2 | ACSF2 | Lys599Asn |
| | CDC23 | Val177Ala |
| | GPC2 | Arg147Trp |
| | **HSD17B11** | Val16Asp |
| | **KCTD5** | Glu196* |
| | MAST2 | Gly1658Trp |
| | MYDGF | Gly22Arg |
| | SIPA1L3 | Gln1126Lys |
| | WWTR1 | Asn213Tyr |
| A4 | AMOTL1 | Gly287Ser |
| | ANKHD1 | Arg192Trp |
| | BAHCC1 | Gln2586Lys |
| | COL6A2 | Glu838* |
| | ETV4 | Lys139Met |
| | **HSD17B11** | Ser172Leu |
| | MGAT5 | Lys554Asn |
| | NUDCD1 | Glu478Lys |
| | TMEM192 | Asp80Gly |
| | TMEM63C | Thr251Lys |
| A5 | AHDC1 | Gln882Leu |
| | COA3 | Gly7Cys |
| | DGKD | Arg590Ser |
| | FBN2 | Thr1526Ala |
| | **KCTD5** | Glu196* |
| | KRIT1 | Gly716Cys |
| | ZNF823 | Gln148* |

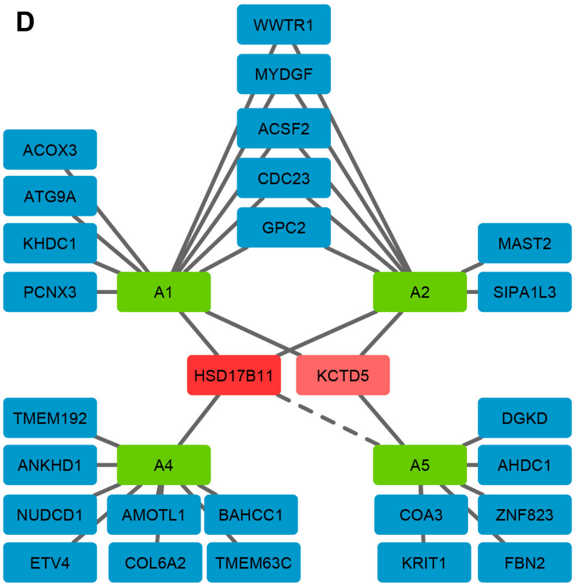

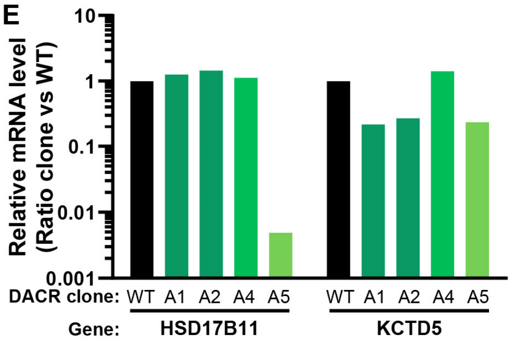

**Supplementary Figure 2: Characterization of DACR clones. A.** Cell viability analysis of DAC-resistant clones (DACR) or wild-type HAP-1 treated for 72 h with indicated concentrations of the proteasome inhibitor bortezomib. **B.** Cell viability analysis of DAC-resistant clones (DACR) or wild-type HAP-1 treated for 72 h with indicated concentrations of the DNA damaging agent doxorubicin. **C.** List of genes carrying near homozygous non- or mis-sense mutations in each clone. For each gene, the impact on the protein sequence is specified. **D.** Graphical representation of the genes identified as mutated in each clone. The genes mutated in more than two clones are highlighted in red. The dashed line indicates that HSD17B11 is not expressed in the DACR#A5 clone. **E.** Histogram representing the ratio between the normalized levels (Reads Per Kilobase Million) of HSD17B11 and KCTD5 mRNAs in each mutant clone as compared to the wild-type HAP-1. Reduced KCTD5 levels in DACR#A1, #A2 and #A5 might result from the KCTD5 mRNA, which carries a premature stop codon, being recognized and processed by non-sense mediated decay.

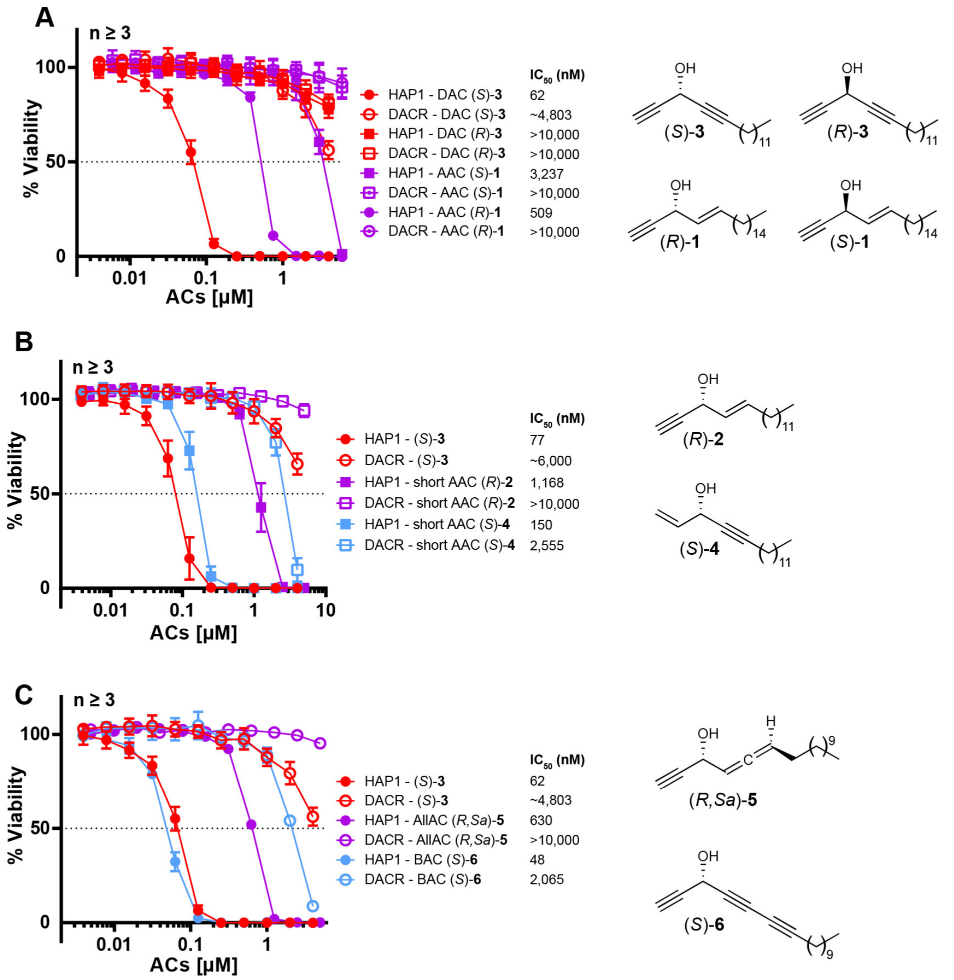

**Supplementary Figure 3: HSD17B11 inactivation confers resistance to multiple alkynylcarbinol-containing molecules (ACs) of similar configuration. A.** Cell viability analysis of DAC-resistant HAP-1 clone #A4 (DACR) treated for 72 h with the indicated concentrations of each of the molecules represented on the right of the panel. **B.** Cell viability analysis of DAC-resistant HAP-1 clone #A4 (DACR) treated for 72 h with the indicated concentrations of each of the molecules represented on the right of the panel. **C.** Cell viability analysis of DAC-resistant HAP-1 clone #A4 (DACR) treated for 72 h with the indicated concentrations of each of the molecules represented on the right of the panel.

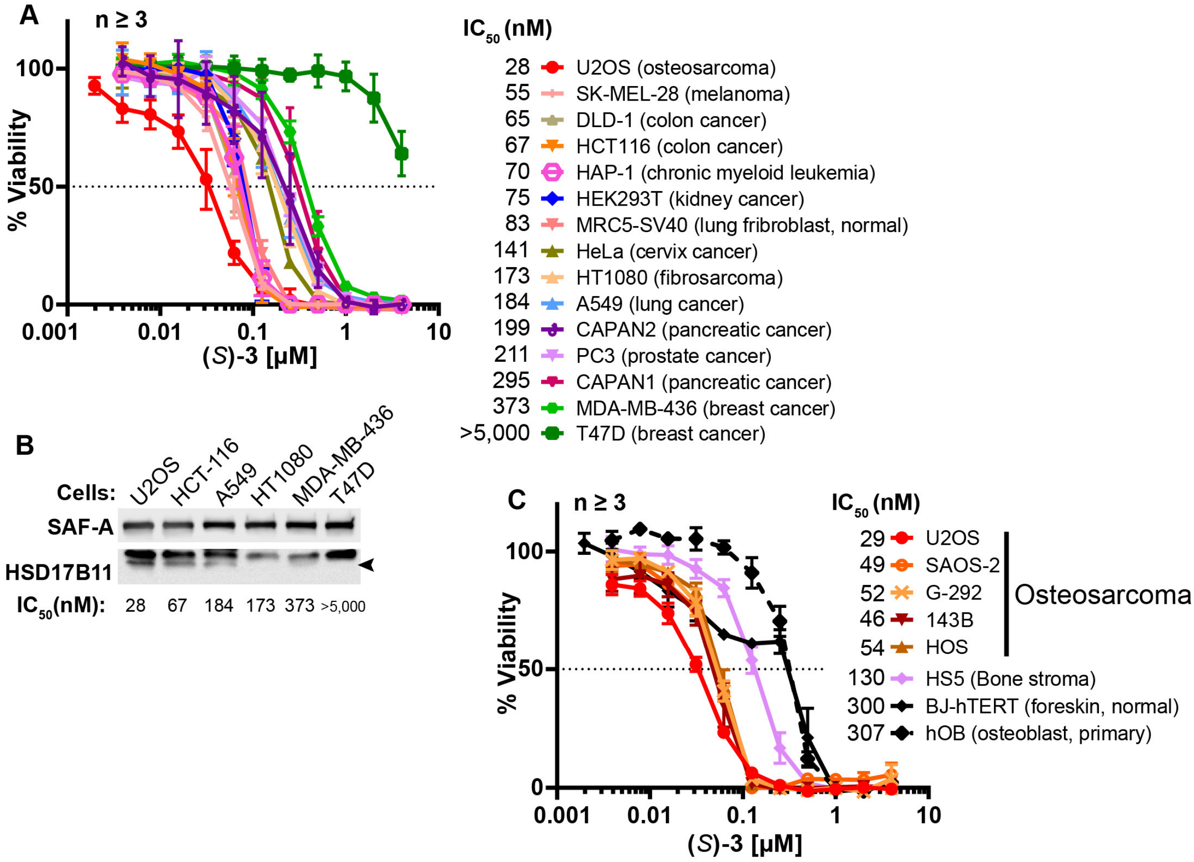

**Supplementary Figure 4: DAC (*S*)-*3* cytotoxic activity is HSD17B11-dependent in multiple cell lines. A.** Cell viability analysis of a panel of 15 cell lines treated for 72 h with DAC (*S*)-**3**. **B.** Analysis of HSD17B11 levels by immunoblotting in 6 selected cell lines from the panel. SAF-A was used as a loading control. **C.** Cell viability analysis of a panel of osteosarcoma cell lines and related control cells treated for 72 h with DAC (*S*)-**3**.

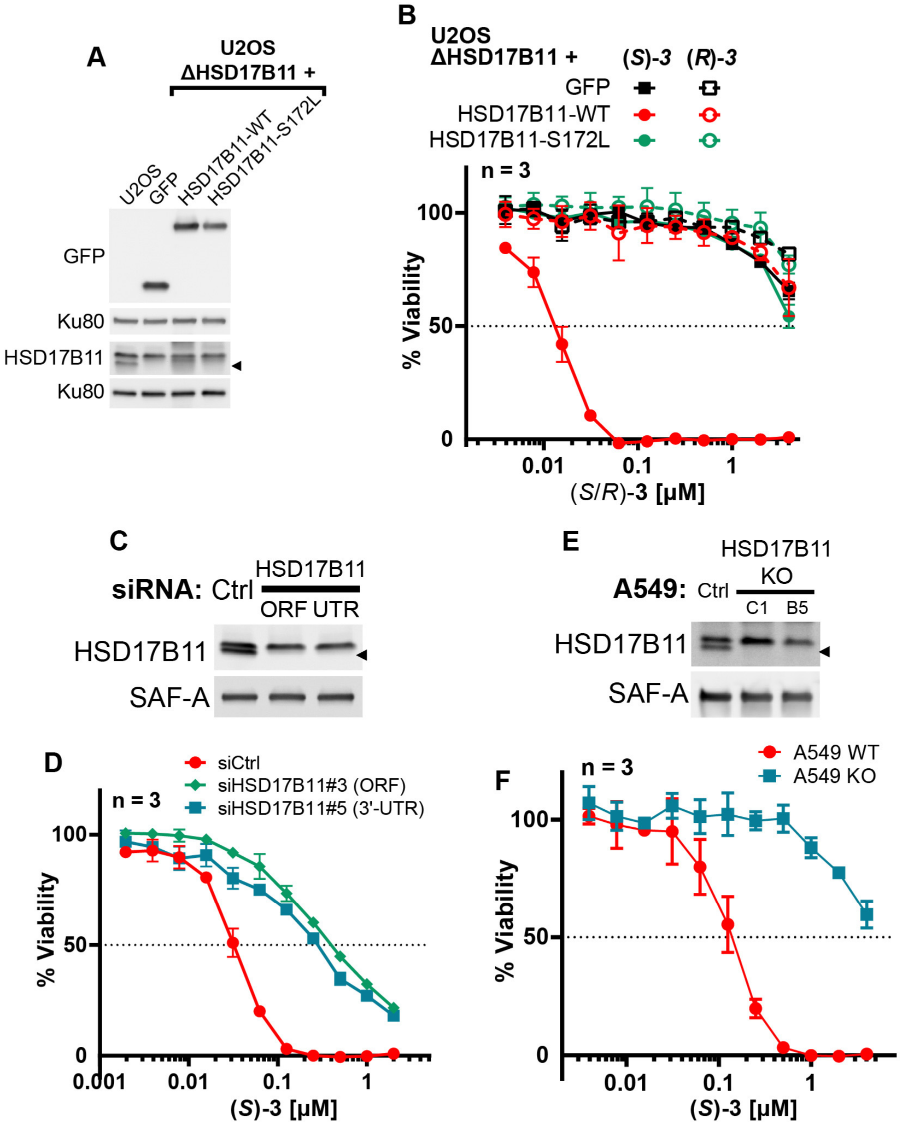

**Supplementary Figure 5: DAC (*S*)-*3* cytotoxic activity is HSD17B11-dependent in multiple cell lines. A.** Analysis of HSD17B11 and HSD17B11-GFP levels in wild-type U2OS or U2OS inactivated for HSD17B11 using CRISPR/Cas9 and stably complemented with GFP or GFP-tagged wild-type or S172L HSD17B11. The black arrow indicates the position of endogenous HSD17B11. Ku80 was used as a loading control. **B.** Cell viability analysis of U2OS inactivated for HSD17B11, complemented as shown in **a** and treated for 72 h with DAC (*S*)- or (*R*)-**3**. **C.** Analysis by immunoblotting of HSD17B11 levels in U2OS cells 72 h after transfection by control or HSD17B11 siRNAs. SAF-A was used as a loading control. **D.** Cell viability analysis of U2OS cells transfected by siRNA as shown in **C** and treated for 72 h with DAC (*S*)-***3***. **E.** Analysis by immunoblotting of HSD17B11 levels in wild-type A549 cells or in individual clones inactivated for HSD17B11 using CRISPR/Cas9. SAF-A was used as a loading control. **F.** Cell viability analysis of wild-type or HSD17B11-deficient (clone #C1) A549 cells treated for 72 h with DAC (*S*)-**3**.

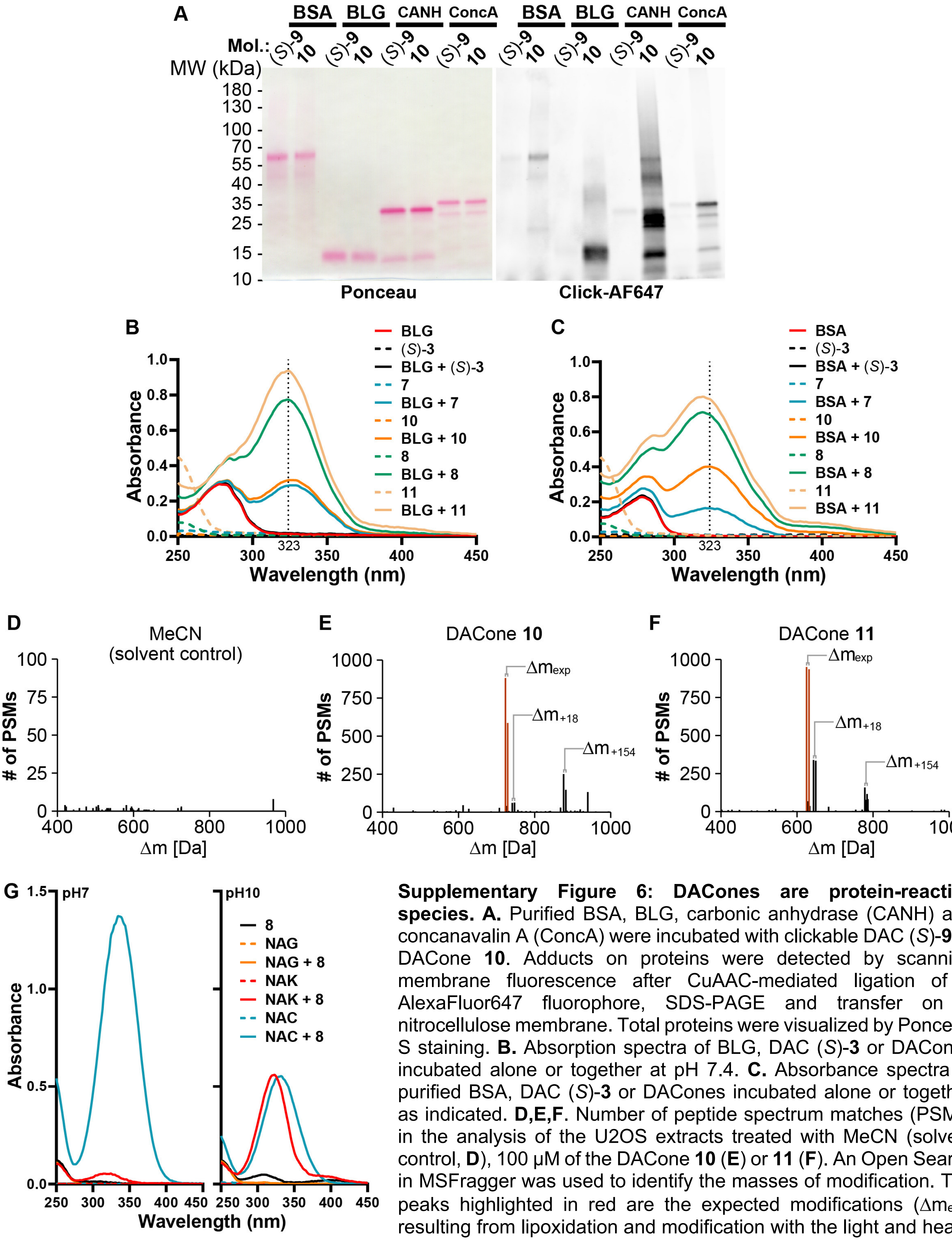

**Supplementary Figure 6: DACones are protein-reactive species. A.** Purified BSA, BLG, carbonic anhydrase (CANH) and concanavalin A (ConcA) were incubated with clickable DAC (*S*)-**9** or DACone **10**. Adducts on proteins were detected by scanning membrane fluorescence after CuAAC-mediated ligation of a AlexaFluor647 fluorophore, SDS-PAGE and transfer on a nitrocellulose membrane. Total proteins were visualized by Ponceau S staining. **B.** Absorption spectra of BLG, DAC (*S*)-**3** or DACones incubated alone or together at pH 7.4. **C.** Absorbance spectra of purified BSA, DAC (*S*)-**3** or DACones incubated alone or together as indicated. **D,E,F**. Number of peptide spectrum matches (PSMs) in the analysis of the U2OS extracts treated with MeCN (solvent control, **D**), 100 µM of the DACone **10** (**E**) or **11** (**F**). An Open Search in MSFragger was used to identify the masses of modification. The peaks highlighted in red are the expected modifications ($\Delta m_{exp}$) resulting from lipoxidation and modification with the light and heavy isoDTB tags, respectively. Further modification of the lipoxidated peptides by reaction with water (+18) or DTT (+154) are also indicated. **G.** Absorption spectra of *N*$_{\alpha}$-acetyl-lysine (NAK), *N*-acetyl-cysteine (NAC), *N*-acetyl-glycine (NAG) or DACone **8** incubated alone or together at pH 7 or at pH 10.

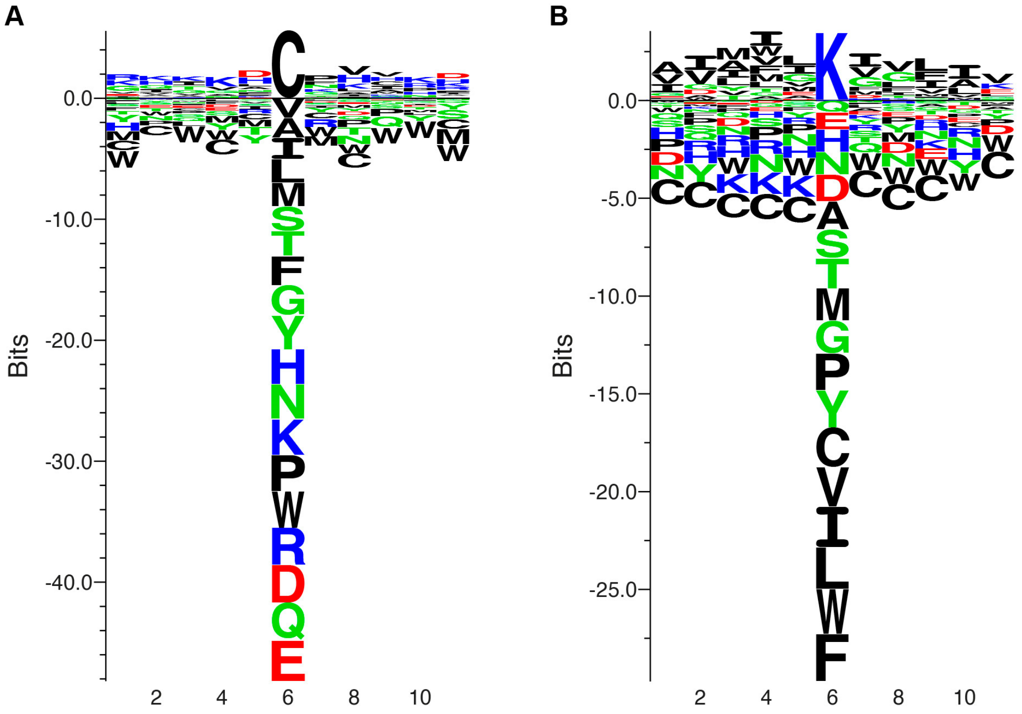

**Supplementary Figure 7: Sequence context of modified cysteines and lysines. A,B.** A 11 amino acids window centered on the cysteines (**A**) or lysines (**B**) modified by the DACone **10** was used to generate a sequence logo. This highlighted that the DACone-modified lysines are preferentially surrounded by hydrophobic amino acids, while no clear enrichment could be identified for the modified cysteines.

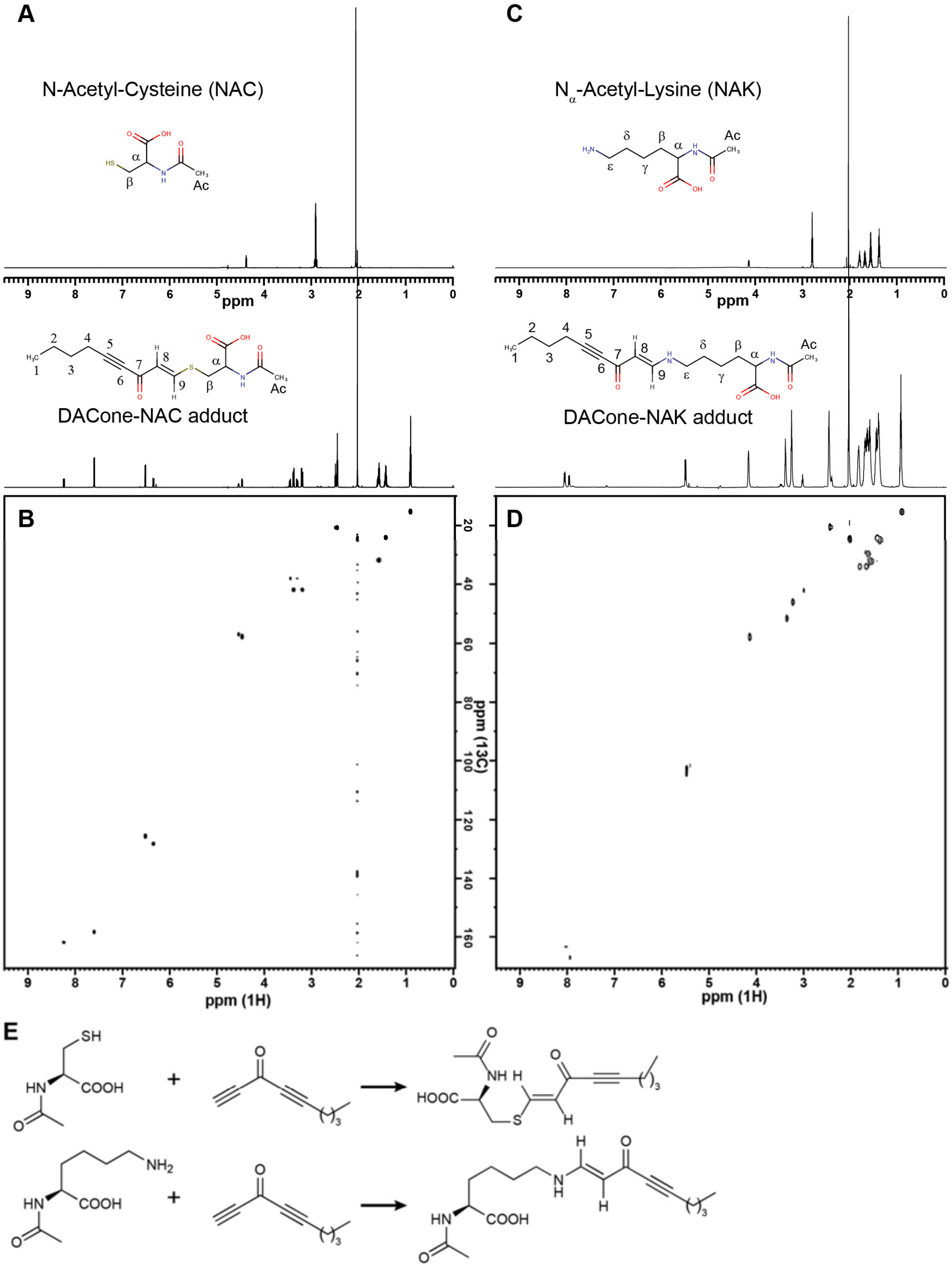

**Supplementary Figure 8: Characterization of DACone reaction products with NAC & NAK. A.** $^{1}$H NMR spectra of DACone **8**-NAC adduct (lower spectrum) or NAC alone (upper spectrum). **B.** $^{1}$H-$^{13}$C HSQC spectra of DACone **8**-NAC adduct. **C.** $^{1}$H NMR spectra of DACone **8**-NAK adduct (lower spectrum) or NAK alone (upper spectrum). **D.** $^{1}$H-$^{13}$C HSQC spectra of DACone **8**-NAK adduct. **E.** Proposed reaction of NAC or NAK with the short DACone **8**.

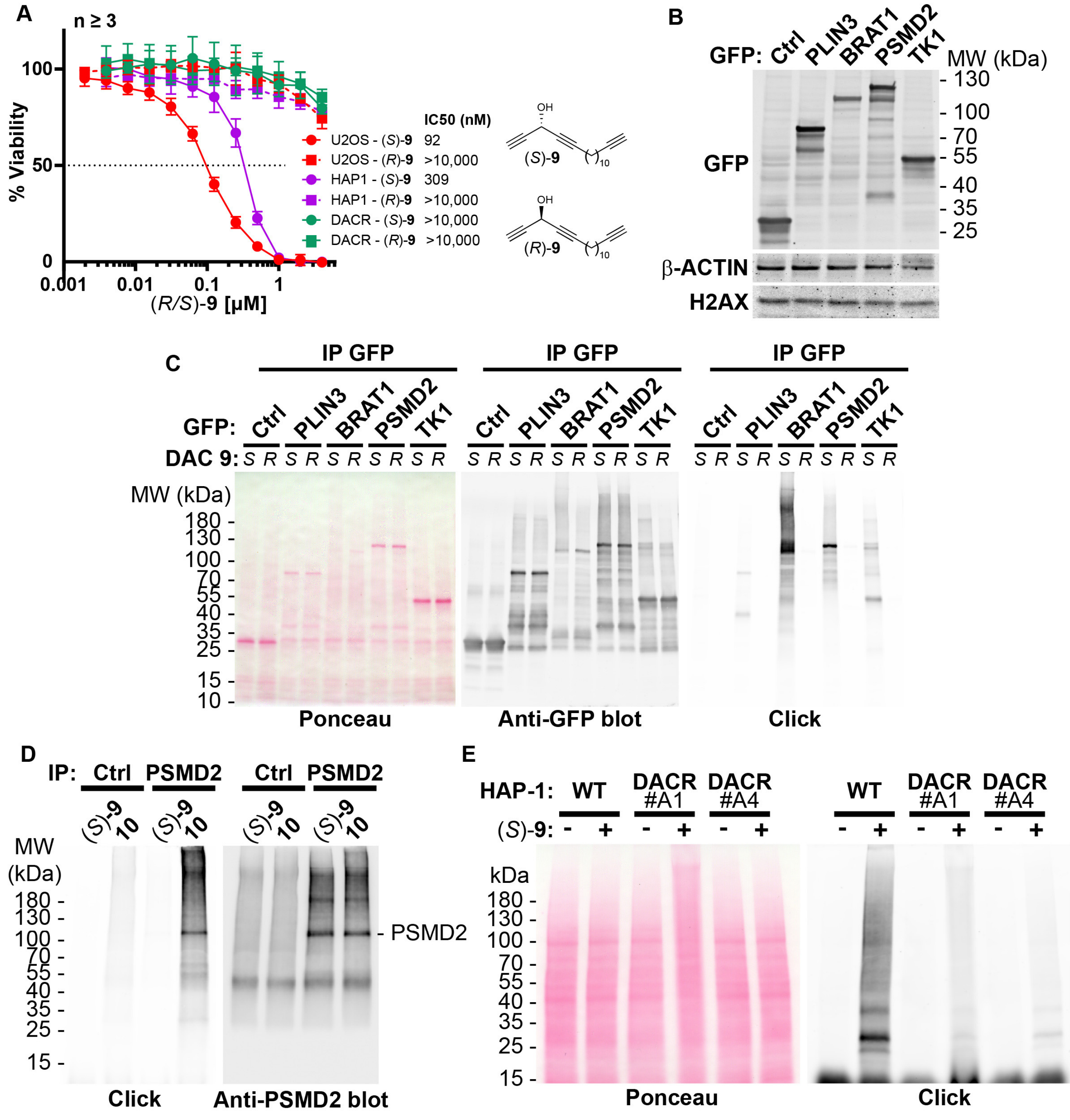

**Supplementary Figure 9: DAC (*S*)-9 is bioactivated into protein-reactive species. A.** Cell viability analysis of U2OS, wild-type HAP-1 or DAC resistant HAP-1 clone A4 treated for 72h with clickable DAC (*S*)-**9** or (*R*)-**9**. **B.** Analysis by immunoblotting of GFP levels in U2OS stably expressing GFP (Ctrl), and GFP-tagged PLIN3, BRAT1, PSMD2 and TK1. **C.** U2OS depicted in B were incubated for 2 h with 2 µM (*S*)-**9** or (*R*)-**9**, proteins were extracted and DAC-modified proteins were detected by CuAAC-mediated ligation of azido-AlexaFluor-647 to clickable molecules, separation by SDS-PAGE, transfer to a membrane which was scanned for fluorescence. **D.** PSMD2 or control immunoprecipitations were performed from extracts of U2OS cells and beads were treated with (*S*)-**9** or DACone **10**. After washes, DAC/DACone-modified proteins were detected by CuAAC-mediated ligation of azido-AlexaFluor-647 to clickable molecules, separation by SDS-PAGE, transfer and scanning membrane fluorescence. PSMD2 was subsequently visualized by immunoblotting. **E.** Wild-type or (*S*)-DAC-resistant HAP-1, clone A1 or A4, cells were untreated or treated 2 h with 2 µM clickable DAC (*S*)-**9**. DAC-modified proteins were detected by CuAAC-mediated ligation of azido-AlexaFluor-647 to clickable molecules, separation by SDS-PAGE, transfer to a membrane which was scanned for fluorescence. PSMD2 was subsequently visualized by immunoblotting.

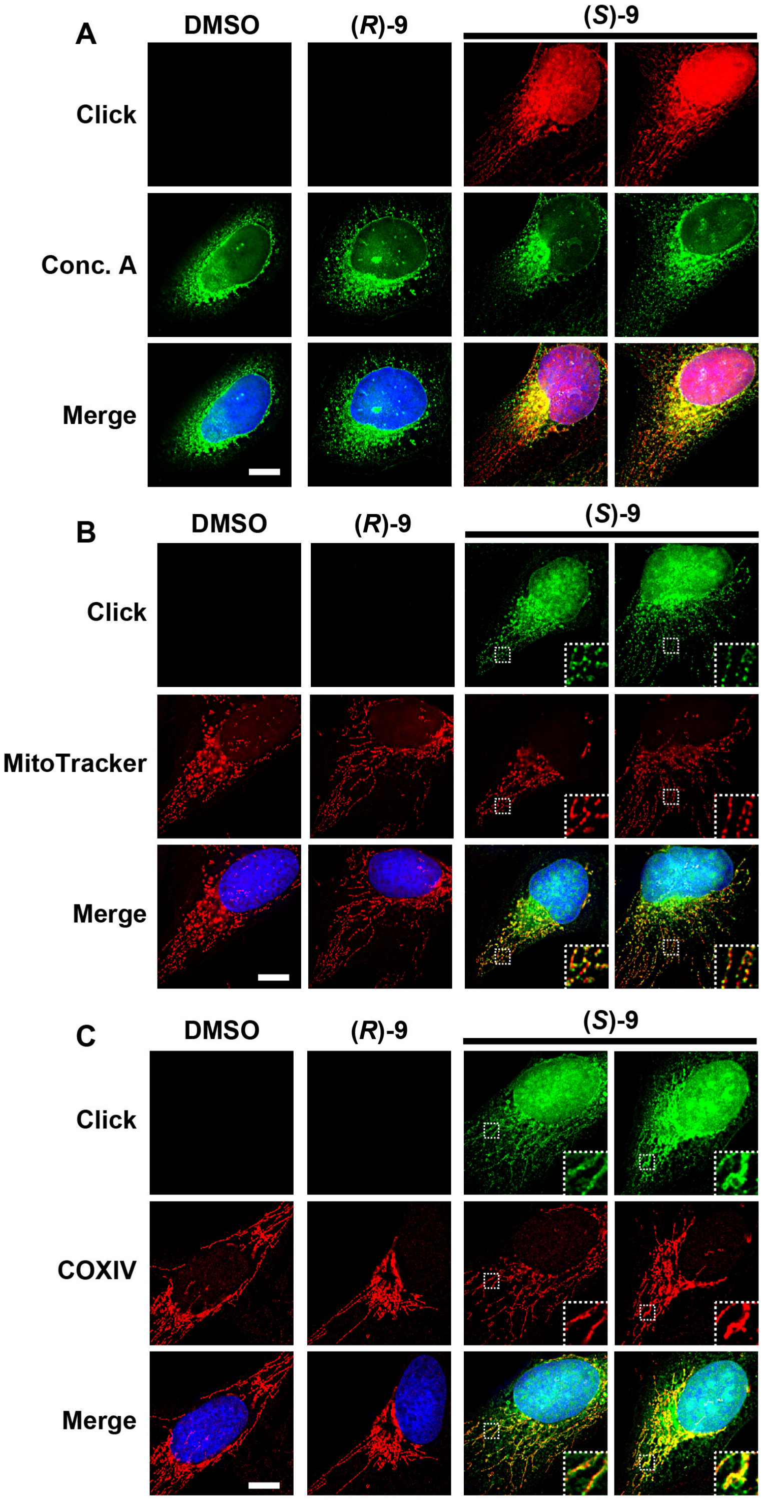

**Supplementary Figure 10: The clickable DAC (*S*)-9 staining colocalizes with the nucleus, ER and mitochondria. A-C.** U2OS cells were treated 2 h with 0.5 μM of DAC (*S*/*R*)-**9**. **A.** After fixation, CuAAC was used to label the DACs with AlexaFluor594-azido, while the ER membranes were labelled using Concanavalin A-AlexaFluor488 (Conc. A). **B.** 30 min before the end of DAC treatment, the cells were incubated with MitoTracker Red CMXRos to label mitochondria. After fixation, click chemistry was used to label clickable DACs with AlexaFluor488-azido. **C.** After fixation, CuAAC was used to label clickable DACs with AlexaFluor488-azido while immunofluorescence with an anti-COXIV antibody coupled with AlexaFluor594 was used to label mitochondria. On all pictures, DAPI was used to stain DNA. White scale bar = 10 μm.

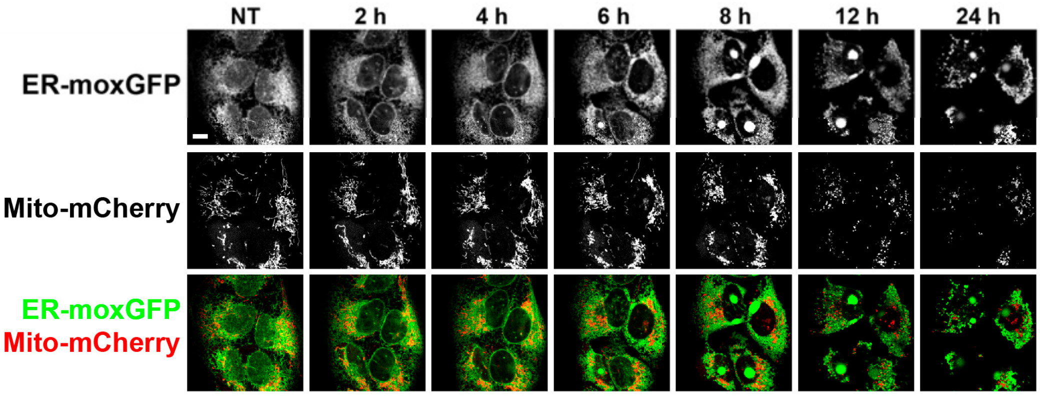

**Supplementary Figure 11: The DAC (*S*)-3 triggers ER-swelling and mitochondrial fission.** U2OS stably co-expressing a GFP variant addressed and retained in the endoplasmic reticulum and mCherry addressed to mitochondria were treated with 1 µM (*S*)-3 and monitored by live imaging.

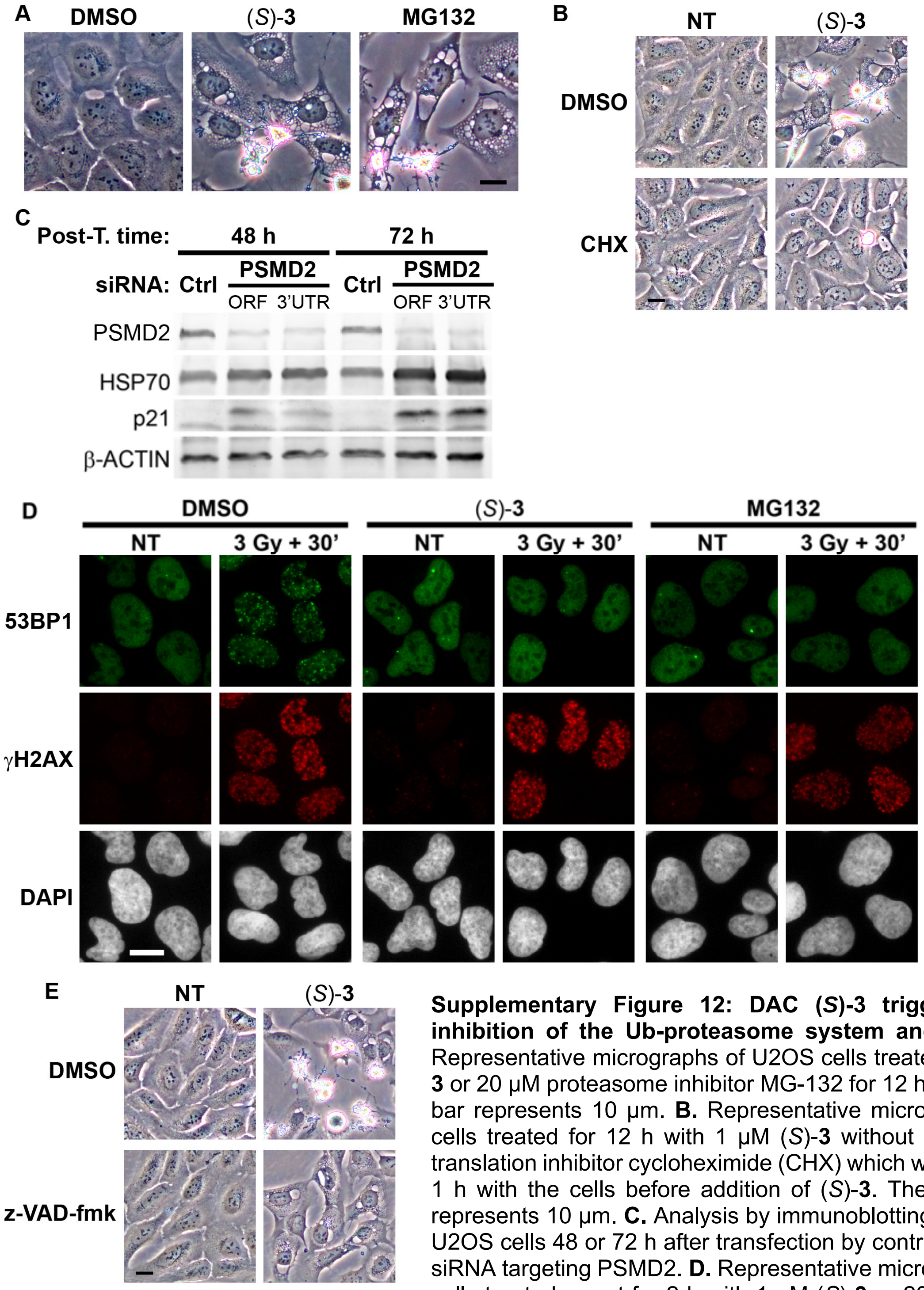

**Supplementary Figure 12: DAC (*S*)-3 triggers ER-stress, inhibition of the Ub-proteasome system and apoptosis. A.** Representative micrographs of U2OS cells treated with 1 µM (*S*)-**3** or 20 µM proteasome inhibitor MG-132 for 12 h. The black scale bar represents 10 µm. **B.** Representative micrographs of U2OS cells treated for 12 h with 1 µM (*S*)-**3** without or with 20 µg/ml translation inhibitor cycloheximide (CHX) which was pre-incubated 1 h with the cells before addition of (*S*)-**3**. The black scale bar represents 10 µm. **C.** Analysis by immunoblotting of extracts from U2OS cells 48 or 72 h after transfection by control siRNA (Ctrl) or siRNA targeting PSMD2. **D.** Representative micrographs of U2OS cells treated or not for 2 h with 1 µM (*S*)-**3** or 20 µM MG132 and irradiated or not with 3 Gy of X-rays, post-incubated for 30 min, fixed and stained for the DNA damage markers gammaH2AX and the 53BP1 protein. Assembly of 53BP1 into foci relies on active ubiquitination at sites of DNA damage. The white scale bar represents 10 µm. **E.** Representative micrographs of U2OS cells treated or not with 1 µM (*S*)-**3** with or without 50 µM z-VAD-fmk. The black scale bar represents 10 µm.

**A**

| Clone | Gene | Mutation |
|---|---|---|
| A4 | CCDC150 | p.Glu413Asp |
| | CCT7 | p.Phe488Ile |
| | FAF2 | p.Glu197Gly |
| | GTSE1 | p.Ser337Asn |
| | HNRNPR | p.Val355Glu |
| | MACROD1 | p.Gly9Cys |
| | MBD2 | p.Arg273* |
| | **RDH11** | p.Arg25Thr |
| | SLC25A38 | p.Phe178Ser |
| | TNK2 | p.Ser808Cys |
| A6 | | |
| B1 | ATRIP | p.Gln483Arg |
| | GTF2I | p.Glu36Gln |
| | POT1 | p.Gly319Glu |
| | **RDH11** | p.Thr226Lys |
| | TMEM132E | p.Pro203Thr |
| | TMEM132E | p.Pro203Arg |
| | TMEM251 | p.Thr21Lys |
| B4 | BCAS3 | p.Gly699Ala |
| | FUZ | p.Trp194* |
| | MGAT4A | p.Tyr269Asp |
| | PBK | p.Leu193Gln |
| | SLC26A6 | p.Leu380Arg |
| | SNX15 | p.Gln4* |
| | UGGT1 | p.Gly746Arg |
| B5 | CAMTA2 | p.Pro184Leu |
| | ITGA7 | p.Gln122* |
| | MTRF1 | p.Gln212* |
| | NCBP1 | p.Gly488Arg |
| | **RDH11** | p.Leu211Pro |
| B6 | CARMIL2 | p.Val251Met |
| | **RDH11** | p.Leu156Gln |
| | SLC31A1 | p.Ile115Thr |
| C1 | ACAD11 | p.Leu78Ile |
| | CAND1 | p.Leu169Arg |
| | NRDE2 | p.Thr280Ile |
| | NUP160 | p.Arg1136Leu |
| | **RDH11** | p.Trp291* |
| | RWDD2B | p.Arg244Thr |

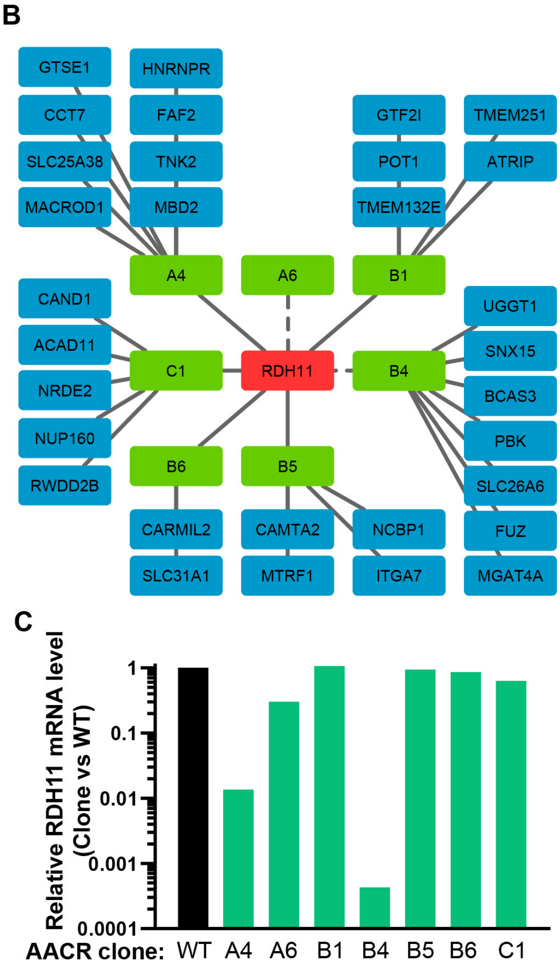

**Supplementary Figure 13: RDH11 is mutated or underexpressed in all AACR clones. A.** List of genes carrying near homozygous non- or mis-sense mutations in each of the seven AACR clones analyzed by RNA-seq. For each gene, the impact on the protein sequence is specified. **B.** Graphical representation of the genes identified as mutated in each clone. The only gene mutated in more than four clones is highlighted in red. The dashed lines indicate that RDH11 is not expressed in the AACR#B4 clone and carries two heterozygous mutations, T227K and R108* in clone AACR#A6. **C.** Histogram representing the ratio between the normalized levels (RPKM) of RDH11 mRNA in each mutant clone as compared to the wild-type HAP-1.

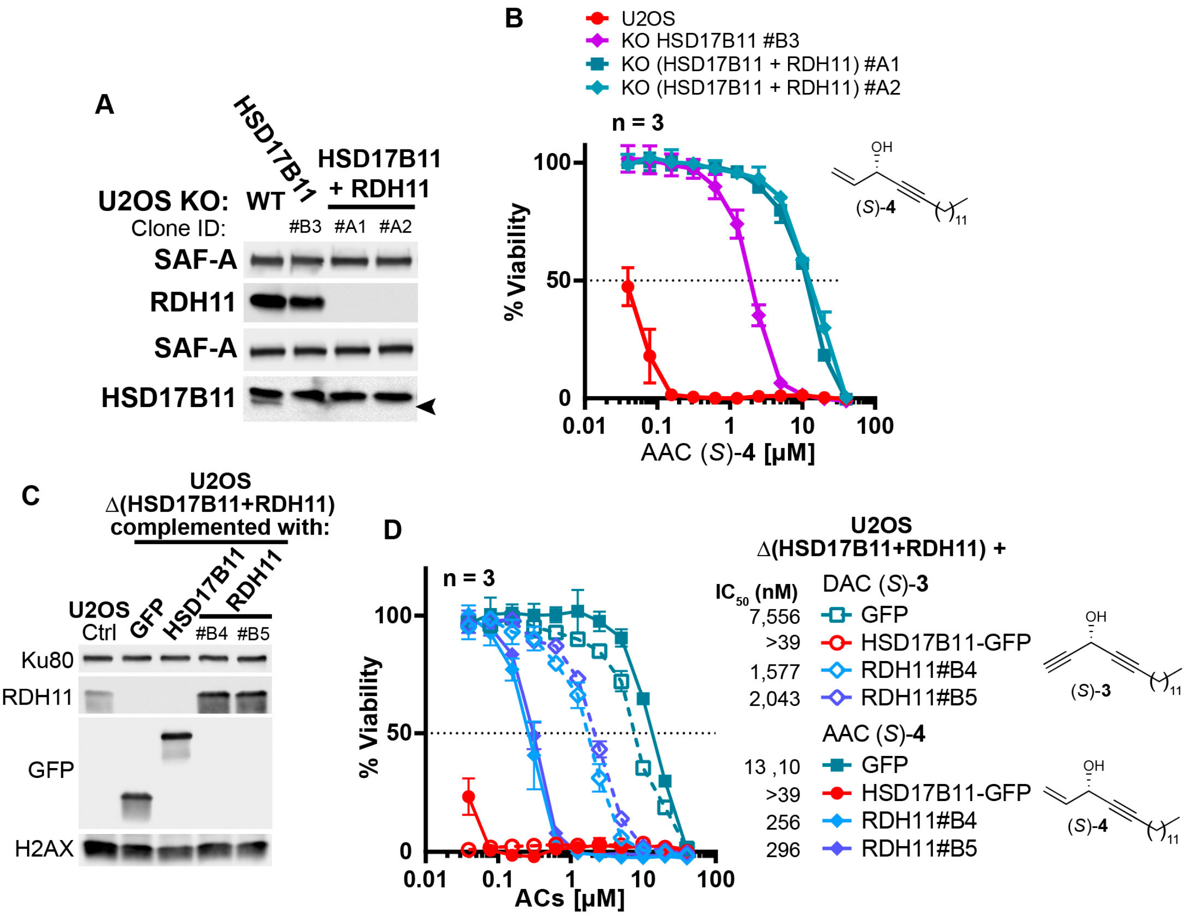

**Supplementary Figure 14: RDH11 bioactivates preferentially *(S)*-AAC over *(S)*-DAC. A.** Analysis by immunoblotting of RDH11 and HSD17B11 levels in wild-type U2OS or U2OS inactivated for HSD17B11 (clone B3) or inactivated for both HSD17B11 and RDH11. SAF-A was used as a loading control. **B.** Cell viability analysis of U2OS either wild-type, inactivated for HSD17B11 or for both HSD17B11 and RDH11, as described in **A**, treated with AAC (*S*)-**4** for 72 h. **C.** Analysis by immunoblotting of RDH11 levels in U2OS either wild-type or inactivated for both HSD17B11 and RDH11 using CRISPR/Cas9 and stably complemented with GFP, HSD17B11-GFP or untagged RDH11 (two different clones). Ku80 and total H2AX were used as loading control. **D.** Cell viability analysis of U2OS inactivated for HSD17B11 and RDH11, stably complemented as described in **C** and treated with DAC (S)-**3** or AAC (*S*)-**4**.

### Supplementary note 1. Strengths of the implemented genomic approach.

Our study highlights the strength of our genomic approach relying on chemical mutagenesis of human haploid cells, selection with a toxic concentration and  the use of RNA-seq to identify simultaneously changes in expression levels and mutations that could confer resistance to cytotoxic drugs. RNA-seq analysis of only four DAC (*S*)-**3**-resistant clones was sufficient to identify HSD17B11 as the DAC (*S*)-**3**-activating enzyme. Three clones carried non-sense mutations, including one on a key catalytic residue (S172), while the other one showed lack of HSD17B11 expression.

The strength of the approach is also reminiscent of the use of chemical mutagenesis. Indeed, while selection of drug-resistant clones based on the appearance of spontaneous mutations is frequently used in bacteria to identify the mechanism of action of small molecules, exploiting their high rate of mutations, human cells are less prone to spontaneous mutations. In human cells, screens based on spontaneous mutations most frequently use DNA repair (mismatch)-deficient cells, such as HCT-116, to increase the mutation rate (see for example (71) and (12)). However, spontaneous mutations can be diverse in nature and in that regard, the use of chemical mutagenesis with ethyl methanesulfonate (EMS), in addition to increasing the rate of resistant clones formation, simplifies the identification of the mutations of interest. Indeed, EMS does not induce deletions or insertion, but principally single nucleotide changes, most frequently transitions (14) which can be selected during the analysis, restricting the number of false positives.

Another strength of this approach is the possibility to readily perform multiple rounds of screening to decipher primary and secondary mechanisms of action for a drug. Here, the DAC (*S*)-**3**-resistant clones could be mutagenized and selected a second time to identify a second SDR, RDH11, as the AAC (*S*)-**4**-bioactivating enzyme. It is noteworthy that the AAC (*S*)-**4** still shows some remaining toxicity on the DAC+AAC-resistant clones (HSD17B11 mutation+RDH11 mutation), which indicates that a third round of mutagenesis, selection and RNA-seq would most likely lead to identify another dehydrogenase, probably in the SDR superfamily, as responsible for the remaining bioactivation. This also illustrates how alkynylcarbinols could constitute a rich reservoir of SDR-specific prodrugs, whose selectivity can be improved through parallel structure-activity relationship studies.

Finally, this approach is remarkably easier and cheaper to implement than loss of function CRIPSR/Cas9 screens and can lead to directly identify essential genes as mediators of drug cytotoxic effect. We recently exemplified this by using this approach to directly identify the DNA topoisomerase II alpha (coded by the essential *TOP2A* gene) as the main driver of the cytotoxic effects of CX-5461, a G-quadruplex ligand and inhibitor of RNA polymerase I (13). The identified non-sense mutations on *TOP2A* dissociate its essential function from its function in generating DNA double-strand breaks upon G-quadruplex stabilization by CX-5461.

### Supplementary note 2. Relevance of the identified mechanism to natural small molecules.

Hundreds of cytotoxic natural compounds have one or several alkynylcarbinol motifs, therefore the mechanism of action identified here could be shared in its principles by these cytotoxic molecules. This is supported by the fact that HSD17B11 was recently identified as mediating the toxicity of the natural compound dehydrofalcarindiol (21), produced in several plants. It is also noteworthy that two other related lipidic natural products, falcarinol and callyspongynic acid, isolated respectively from several *Apiaceae* plant species and the marine sponge *Callyspongia truncata*, have been identified as covalently binding to cellular proteins (72,73), suggesting that they could also be bioactivated into protein-reactive species by a yet unidentified mechanism. Consequently, we propose that alkynylcarbinol-containing natural molecules represent a defense mechanism through bioactivation by specific SDRs in the body of predators, pathogens or parasites. In agreement, fulvindione, the AAC-oxidized form of the cytotoxic *Haliclona fulva*-produced fulvinol, was found in the body of the dorid nudibranch *Peltodoris atromaculata* feeding on *Haliclona* (74). The presence of multiple alkynylcarbinol motifs in some of these natural compounds could provide modular pro-cytotoxic agents bioactivable by SDRs in different organisms. The fact that HSD17B11 mediates the toxicity of several natural compounds of unrelated origins questions the reasons for it being the common bioactivating enzyme of these prodrugs. HSD17B11 is localized at the endoplasmic reticulum and on lipids droplets (16). Strikingly, lasonolide A, a macrolide isolated from the marine sponge *Forcepia sp.*, was also identified as a prodrug bioactivated by a ER/lipid-droplet resident enzyme, the serine hydrolase LDAH (75). Lipid droplets are conserved organelles (76) and multiple lipidic natural compounds might exploit lipid droplets-associated enzymes for cytotoxicity.

### Supplementary note 3. Parallel between DAC and calicheamicin.

Another family of alkyne-containing natural small molecules with potent cytotoxic activity which was harnessed to develop new anticancer treatment is the enediyne family, with its most known representative, calicheamicin-gamma1 (cali), extracted from the soil bacteria *Micromonospora echinospora* (77). Cali shows a cytotoxic activity in the picomolar range that is exploited by two anticancer treatments, gemtuzumab ozogamicine (Mylotarg®) and inotuzumab ozogamicin (Besponsa®), consisting respectively of a CD33-cali or CD22-cali antibody-drug conjugates used for the treatment of acute myeloid leukemia (78). In the enediyne motif, the equivalent position of the DAC carbinol unit >CHOH is replaced by an ethylene unit -CH=CH-. Cali is also a prodrug which is activated by the intracellular reducing environment triggering the reduction, mainly carried by cellular glutathione, of a trisulfide unit to a thiol group, initiating a cascade process by an intramolecular Michael addition leading to the DNA-damaging biradical species generated by Bergman cyclization. This, together with the binding of cali to the minor groove of DNA through its aryloligosaccharide moiety, allows the reaction of cali with both DNA strands by promoting hydrogen abstraction on the deoxyribose part of DNA, leading to cytotoxic DNA double-strand breaks (79).

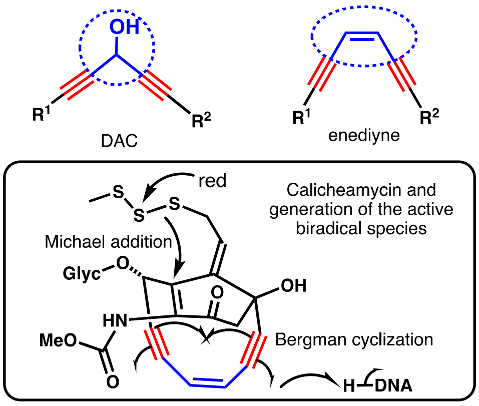

### Supplementary note 4. Synthesis of novel compounds.

All reagents were obtained from commercial suppliers and used without any further purification. Reactions were run under nitrogen or argon atmosphere in oven-dried glassware. Standard inert atmosphere techniques were used in handling air and moisture sensitive reagents. Dichloromethane ($CH_2Cl_2$) and tetrahydrofuran (THF) were obtained by filtration through a drying column on a filtration system. Thin-layer chromatography analyses were performed on precoated, aluminum-backed silica gel (Merck 60 F254). Visualization of the developed chromatogram was performed by UV light (254 nm) and using aqueous potassium permanganate ($KMnO_4$) stain. Flash column chromatography was performed using flash silica gel (SDS 35-70 mm or 60 Å, C.C 70-200 µm). Nuclear magnetic resonance spectra were recorded on Bruker Advance 300 or 400 MHz spectrometers. Chemical shifts for $^1$H NMR spectra are quoted in parts per million relative to residual solvent peak. Data are reported as follows: chemical shift, multiplicity (s = singlet, d = doublet, t = triplet, q = quartet, qn = quintet, m = multiplet), coupling constant in Hz and integration. Chemical shifts for $^{13}$C NMR spectra are quoted in parts per million spectra relative to residual solvent peak. All $^{13}$C NMR spectra were obtained with complete proton decoupling. Infrared analyses were run on a Perkin-Elmer Spectrum 100 FT-IR spectrometer or a Thermo-Nicolet Diamond ATR (4 cm$^{-1}$ of resolution, 16 scans) equipped with a DTGS detector and are reported in reciprocal centimeters (cm$^{-1}$). High-resolution mass spectrometry (HRMS) was performed on a Thermo-Finnigan MAT 95 XL instrument.

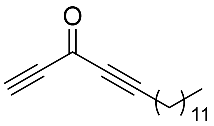

**heptadeca-1,4-diyn-3-one (7)**: To a solution of the previously described racemic dialkynylcarbinol ***3*** (42.5 mg, 0.17 mmol) in $CH_2Cl_2$ (4.0 mL) was added Dess-Martin periodinane (109 mg, 0.26 mmol, 1.5 eq) at RT. The reaction mixture was stirred until completion (TLC monitoring). After 1.5 h, water (2 mL) was slowly added to the solution. Layers were separated and the aqueous layer was extracted three times with $CH_2Cl_2$. The combined organic layers were dried over $MgSO_4$ and concentrated under reduced pressure at room temperature. Flash chromatography on silica (gradient elution up to 10% $Et_2O$ in pentane) of the crude mixture afforded DACone ***7*** (42 mg, 0.17 mmol, quant. yield) as a slightly yellow oil. **$^1$H NMR** (300 MHz, $CD_3CN$) δ 3.79 (s, 1H), 2.45 (t, $J$ = 7.0 Hz, 2H), 1.65 – 1.50 (m, 2H), 1.45 – 1.20 (m, 18H), 0.88 (t, $J$ = 6.7 Hz, 3H). **$^{13}$C NMR** (75 MHz, $CD_3CN$) δ: 161.2, 98.2, 82.8, 82.4, 80.3, 32.6, 30.3, 30.2, 30.1, 30.0, 29.6, 29.4, 28.1, 23.4, 19.4, 14.4. **HRMS-DCI** ($CH_4$): *m/z* calcd for $C_{17}H_{27}O$ $[M]^{+\cdot}$: 247.2062, found: 247.2060. **FTIR** (cm$^{-1}$) (neat): $\nu$ 2925, 2855, 2223, 2098, 1636, 1211. DACone ***7*** was dissolved in $CH_3CN$ at 1 mM.

**Synthesis of nona-1,4-diyn-3-one (8)**

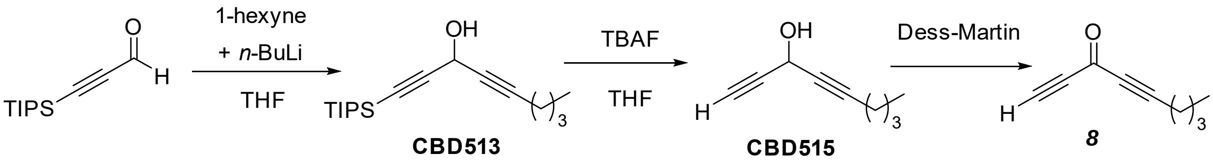

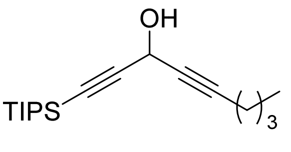

**1-[tris(propan-2-yl)silyl]-nona-1,4-diyn-3-ol (CBD513)**: To a solution of 1-hexyne (0.273 mL, 2.97 mmol, 1 eq.) in THF (5 mL) under stirring at 0 °C was added *n*-butyllithium (2.5 M solution in hexane, 1.19 mL, 2.97 mmol, 1 eq.). The solution was stirred for 10 minutes at 0 °C and then 30 minutes at RT before treatment with 3-[tris(propan-2-yl)silyl]-prop-2-ynal (625 mg, 2.97 mmol, 1 eq.) at 0 °C. The mixture was stirred 10 minutes at 0 °C and then overnight at RT. After treatment with a saturated aqueous $NH_4Cl$ solution, the aqueous layer was extracted three times with $Et_2O$. The combined organic layers were washed with brine, dried over $MgSO_4$ and concentrated under reduced pressure. The residue was purified by silica gel column chromatography (cyclohexane/$Et_2O$ 9:1) to give **CBD513** (480 mg, 1.64 mmol, 55%) as a colorless oil. **$^1$H NMR** (300 MHz, $CDCl_3$) $\delta$ 5.11 (s, 1H), 2.25 (td, *J* = 6.8 Hz, 2.1 Hz, 2H), 1.40-1.54 (m, 4H), 1.1 (s, 21H), 0.93 (t, *J* = 7.2 Hz, 3H). **$^{13}$C NMR** (75 MHz, $CDCl_3$): $\delta$ 104.8, 85.3, 85.1, 77.7, 52.8, 30.3, 21.8, 18.5, 18.3, 13.5, 11.1. **HRMS-DCI** ($CH_4$): *m/z* calcd for $C_{18}H_{33}OSi$ $[M+H]^+$: 293.2290, found: 293.2301.

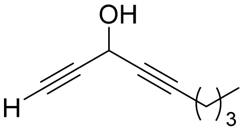

**nona-1,4-diyn-3-ol (CBD515)**: To a solution of alcohol **CBD513** (480 mg, 1.64 mmol) in THF (50 mL) under stirring at 0 °C was added dropwise tetra-*n*-butylammonium fluoride (1 M in THF, 4.92 mL, 4.92 mmol). Then, the mixture was stirred for 90 minutes at room temperature (RT) before addition of a saturated aqueous $NH_4Cl$ solution. The aqueous layer was extracted with $Et_2O$. The combined organic layers were washed with brine, dried over $MgSO_4$ and concentrated under reduced pressure. The residue was purified by silica gel column chromatography (pentane/$Et_2O$ 95:5) to give **CBD515** (180 mg, 80%) as a colorless oil. **$^1$H NMR** (300 MHz, $CDCl_3$) $\delta$ 5.12 (q, *J* = 2.1 Hz, 1H), 2.56 (d, *J* = 2.3 Hz, 1H), 2.25 (td, *J* = 7 Hz, 2.1 Hz, 2H) 1.34-1.64 (m, 4H), 0.9 (t, *J* = 7.2 Hz, 3H). **$^{13}$C NMR** (100 MHz, $CDCl_3$): $\delta$ 96.0, 81.5, 77.0, 72.1, 52.1, 21.9, 18.3, 17.7, 13.5.

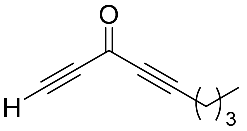

**nona-1,4-diyn-3-one (8):** To a solution of diynol **CBD515** (163 mg, 1.2 mmol) in $CH_2Cl_2$ (30 mL) was added Dess-Martin periodinane (0.3 M in $CH_2Cl_2$, 8 mL, 2.4 mmol) at RT and the resulting mixture was stirred for 4 h. The reaction was quenched by addition of a saturated aqueous $NH_4Cl$ solution. The aqueous layer was extracted with $CH_2Cl_2$. The combined organic layers were washed with brine, dried over $MgSO_4$ and concentrated under reduced pressure. The residue was purified by silica gel column chromatography (pentane/ $Et_2O$ 9:1) to give **8** (75 mg, 46%) as a colorless oil. **$^1$H NMR** (300 MHz, $CDCl_3$) $\delta$ 3.28 (s, 1H), 2.44 (t, *J* = 7 Hz, 2H), 1.55-1.72 (m, 2H), 1.39-1.55 (m, 2H), 0.95 (t, *J* = 7.2 Hz 3H). **$^{13}$C NMR** (75 MHz, $CDCl_3$) $\delta$ 160.4, 96.7, 82.2, 81.9, 77.9, 29.4, 21.9, 18.8, 13.4. **HRMS-DCI** ($CH_4$) *m/z* calcd for $C_9H_{11}O$, $[M+H]^+$: 135.0810, found: 135.0806.

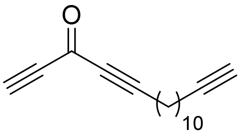

**Synthesis of heptadeca-1,4,16-triyn-3-one (10):** To a solution of the previously described racemic clickable dialkynylcarbinol **9** (30.0 mg, 0.12 mmol) in $CH_2Cl_2$ (2.9 mL) was added Dess-Martin periodinane (78 mg, 0.18 mmol, 1.5 eq) at RT. The reaction mixture was stirred until completion (TLC monitoring). After 80 min, water (1.5 mL) was slowly added to the solution. Layers were separated and the aqueous layer was extracted three times with $CH_2Cl_2$. The combined organic layers were dried over $MgSO_4$ and concentrated under reduced pressure at RT. Flash chromatography on silica (gradient elution up to 15% $Et_2O$ in pentane) of the crude mixture afforded clickable DACone **10 (**26.5 mg, 0.12 mmol, 89% yield) as a colorless oil. **$^1$H NMR** (300 MHz, $CD_3CN$) $\delta$ 3.80 (s, 1H), 2.45 (t, *J* = 7.0 Hz, 2H), 2.20 – 2.10 (m, 3H), 1.65 – 1.52 (m, 2H), 1.52 – 1.25 (m, 14H). **$^{13}$C NMR** (75 MHz, $CD_3CN$) $\delta$ 161.2, 98.2, 85.5, 82.8, 82.4, 80.3, 69.5, 30.1, 30.0, 29.7, 29.6, 29.4, 29.4, 29.3, 28.1, 19.4, 18.7. **HRMS-DCI** ($CH_4$): *m/z* calcd for $C_{17}H_{23}O$ $[M+H]^+$: 243.1749, found: 243.1761. **FTIR** ($cm^{-1}$) (neat): $\nu$ 3288, 2923, 2851, 2214, 2100, 1629, 1207. Clickable DACone **10** was dissolved in $CH_3CN$ at 1 and 20 mM.

**Synthesis of deca-1,4,9-triyn-3-one (11):**

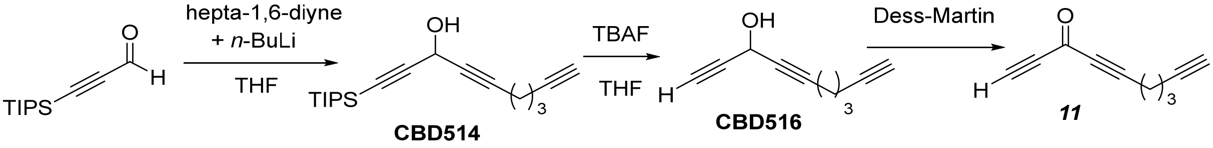

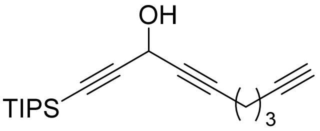

**1-[tris(propan-2-yl)silyl]-deca-1,4,9-triyn-3-ol (CBD514)**: To a solution of hepta-1,6-diyne (0.342 mL, 3 mmol) in THF (5 mL) under stirring at -78 °C was added dropwise n-butyllithium (2.5 M in hexane, 1.2 mL, 3 mmol). The solution was stirred for 1 hour at – 78 °C and 30 minutes at RT before treatment with 3-[tris(propan-2-yl)silyl]-prop-2-ynal (631 mg, 3 mmol). The mixture was allowed to warm slowly up to RT and stirred overnight. After treatment with a saturated aqueous $NH_4Cl$ solution, the aqueous layer was extracted three times with $Et_2O$. The combined organic layers were washed with brine, dried over $MgSO_4$ and concentrated under reduced pressure. The residue was purified by silica gel column chromatography (cyclohexane/$Et_2O$ 99:1) to give **CBD514** (490 mg, 54%) as a colorless oil. **$^1$H NMR** (400 MHz, $CDCl_3$) *d* 5.11 (s, 1H), 2.39 (td, *J* = 7.0, 2.1 Hz, 2H,), 2.33 (td, *J* = 7.0, 2.7 Hz, 2H), 1.97 (t, J = 2.7 Hz, 1H), 1.76 (qn, *J* = 7.0 Hz, 2H), 1.10 (s, 21H). **$^{13}$C NMR** (75 MHz, $CDCl_3$) $\delta$ 104.6, 85.5, 84.0, 83.3, 78.5, 68.9, 52.7, 27.2, 18.5, 17.7, 17.4, 11.1.

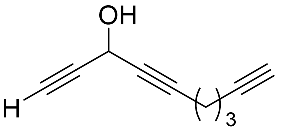

**deca-1,4,9-triyn-3-ol (CBD516)**: To a solution of **CBD514** (490 mg, 1.60 mmol) in THF (50mL) under stirring at 0°C was added dropwise tetra-*n*-butylammonium fluoride (1M in THF, 4.8 mL, 4.8 mmol). Then, the mixture was stirred for 2 h at RT before addition of a saturated aqueous $NH_4Cl$ solution. The aqueous layer was extracted with $Et_2O$. The combined organic layers were washed with brine, dried with $MgSO_4$ and concentrated under reduced pressure. The residue was purified by silica gel column chromatography (pentane/$Et_2O$ 9:1) to give **CBD516** (130 mg, 47%) as a colorless oil. **$^1$H NMR** (300 MHz, $CDCl_3$) $\delta$ 5.12 (q, *J* = 2.1 Hz, 1H), 2.57 (d, *J* = 2.3 Hz, 1H), 2.40 (td, *J* = 7.1, 2.1 Hz, 2H), 2.33 (td, *J* = 7.0, 2.7 Hz, 2H), 1.99 (t, *J* = 2.7 Hz, 1H), 1.68-1.85 (m, 2H). **$^{13}$C NMR** (75 MHz, $CDCl_3$) $\delta$ 84.7, 83.3, 81.3, 77.7, 72.3, 69.0, 52.1, 27.1, 17.7, 17.5. **HRMS-DCI** ($CH_4$) *m/z* calcd for $C_{10}H_8$ $[M-H_2O]^+$: 128.0626, found: 128.0623.

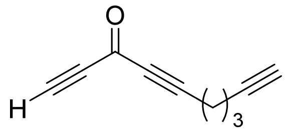

**deca-1,4,9-triyn-3-one (11)**: To a solution of **CBD516** (130 mg, 0.74 mmol) in $CH_2Cl_2$ (30 mL) was added Dess-Martin periodinane (0.3 M in $CH_2Cl_2$, 3.7 mL, 1.12 mmol) at RT and the resulting mixture was stirred for 4 h. The reaction was quenched by addition of a saturated aqueous $NH_4Cl$ solution. The aqueous layer was extracted with $CH_2Cl_2$. The combined organic layers were washed with brine, dried with $MgSO_4$ and concentrated under reduced pressure. The residue was purified by silica gel column chromatography (pentane/$Et_2O$ 9:1) to give **11** (65 mg, 50%) as a colorless oil. **$^1$H NMR** (300 MHz, $CDCl_3$) $\delta$ 3.3 (s, 1H), 2.60 (t, *J* = 7.1 Hz, 2H), 2.37 (td, *J* = 6.9, 2.7 Hz, 2H), 2.02 (t, *J* = 2.6 Hz, 1H), 1.86 (qn, *J* = 7.1 Hz, 2H). **$^{13}$C NMR** (75 MHz, $CDCl_3$) $\delta$ 160.1, 95.2, 82.5, 82.2, 82.1, 78.2, 69.6, 26.3, 18.1, 17.6. **HRMS-DCI** ($CH_4$) *m/z* calcd for $C_{10}H_9O$, $[M+H]^+$: 145.0653, found: 145.0649.

**Synthesis of heptadec-1-en-4-yne-3,6-diol (12):**

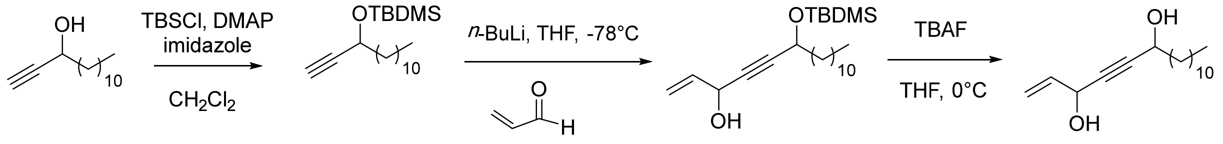

MVB36 MVB39 MVBL05 12

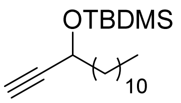

***tert*-butyldimethyl(tetradec-1-yn-3-yloxy)silane (MVB39)**: Imidazole (310 mg, 4.56 mmol), DMAP (19.9 mg, 0.163 mmol,) and TBSCl (737 mg 4.89 mmol) were added to a solution of **MVB36** (80) (3.26 mmol, 686 mg) in anhydrous $CH_2Cl_2$ (10 mL) at RT. After stirring overnight, the reaction mixture was quenched with $H_2O$ (10 mL) and the aqueous layer was extracted with $CH_2Cl_2$ (3 x 10 mL). The combined organic layers were dried over anhydrous $MgSO_4$ and concentrated under reduced pressure. The residue was purified by silica gel column chromatography (petroleum ether) to give *tert*-butyldimethyl(tetradec-1-yn-3-yloxy)silane (655 mg, 62%) as a colourless oil. **$^1$H NMR** (300 MHz, $CDCl_3$) $\delta$ 4.33 (td, *J* = 6.5, 2.1 Hz, 1H), 2.37 (d, *J* = 2.1 Hz, 1H), 1.70 – 1.63 (m, 2H), 1.46 – 1.36 (m, 2H), 1.34 – 1.20 (s, 16H), 0.91 (s, 9H), 0.88 (t, *J* = 6.9 Hz, 3H), 0.13 and 0.11 (2 s, 2 x 3H). **$^{13}$C NMR** (75 MHz, $CDCl_3$) $\delta$ 86.0, 72.0, 62.9, 38.7, 32.1, 29.8, 29.8, 29.7, 29.7, 29.5, 29.4, 25.9 (3 $CH_3$), 25.3, 22.8, 18.4, 14.3, -4.4, -4.9. **HRMS-DCI** ($CH_4$): *m/z* $[M+H]^+$: calcd for $C_{20}H_{41}OSi$: 325.2927 found: 325.2918. **FTIR** ($cm^{-1}$) (neat): $\nu$ 3312, 2924, 2854, 1463, 1250, 1094, 835, 776, 652, 626, 559.

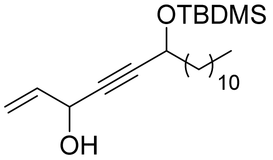

**6-((*tert*-butyldimethylsilyl)oxy)heptadec-1-en-4-yn-3-ol (MVBL05)**: To a solution of **MVB39** (117 mg, 0.361 mmol,) in THF (8 mL) at -78 ºC, *n*-butyllithium solution (2.5 M in hexane, 159 µL, 0.40 mmol) was added dropwise. After 30 min, a solution of acrolein (80.7 mg, 1.44 mmol) in THF (0.5 mL) was added dropwise. The reaction was warmed up to RT and maintained under stirring overnight. It was quenched by the addition of saturated $NH_4Cl$ aqueous solution (5 mL) and extracted with EtOAc (3 × 8 mL). The combined organic layers were dried over anhydrous $MgSO_4$, and concentrated under reduced pressure. The residue was purified by silica gel column chromatography (pentane/EtOAc 25:1) to give 6-((*tert*-butyldimethylsilyl)oxy)heptadec-1-en-4-yn-3-ol (38 mg, 27%). Mixture of diastereoisomers **$^1$H NMR** (400 MHz, $CDCl_3$): $\delta$ 5.96 (ddd, *J* = 17.0, 10.2, 5.3 Hz, 1H), 5.45 (dd, *J* = 17.0, 1.3 Hz, 1H), 5.21 (d, *J* = 10.2 Hz, 1H), 4.95 – 4.85 (m, 1H) 4.39 (td, *J* = 6.5, 1.6 Hz, 1H), 1.75 – 1.60 (m, 2H),

1.47 – 1.35 (m, 2H), 1.35 – 1.25 (m, 16H), 0.90 (s, 9H), 0.85 (t, *J* = 6.9 Hz, 3H), 0.12 and 0.10 (2 s, 2 x 3H). **$^{13}$C NMR** (100 MHz, $CDCl_3$): *δ* 137.15/137.13, 116.48/116.46, 88.59/88.56, 82.64/82.60, 63.4, 63.1, 38.7, 32.1, 29.9, 29.8, 29.8, 29.7, 29.7, 29.5, 29.4, 26.0, 25.4, 22.8, 18.4, 14.3, -4.3, -4.8. **HRMS-DCI** ($CH_4$): *m/z* calcd for $C_{23}H_{45}O_2Si$ [M+H]$^+$: 381.3189, found: 381.3186. **FTIR** (cm$^{-1}$) (neat): *ν* 3343, 2924, 2854, 1463, 1251, 1092, 984, 927, 835, 776.

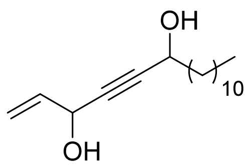

**heptadec-1-en-4-yne-3,6-diol (12)**: To a stirring solution of **MVBL05** (0.10 mmol, 38 mg) in THF (5 mL) at 0 °C, was added a solution of tetrabutylammonium fluoride (1 M in THF, 198 µL, 0.198 mmol). The mixture was stirred for 4 h at RT. The reaction was quenched by addition of saturated aqueous solution of $NH_4Cl$, followed by extraction with EtOAc (3 x 5 mL). The combined organic fractions were dried over anhydrous $MgSO_4$, filtrated and concentrated under reduced pressure. The residue was purified by silica gel column chromatography (pentane/EtOAc 10:3) to give heptadec-1-en-4-yne-3,6-diol (**12**) as a colourless oil (17 mg, 64%). Mixture of diastereoisomers **$^1$H NMR** (400 MHz, $CDCl_3$): *δ* 5.97 (ddd, *J* = 17.0, 10.1, 5.3 Hz, 1H), 5.45 (dt, *J* = 17.0, 1.5 Hz, 1H), 5.23 (dt, *J* = 10.2, 1.3 Hz, 1H), 4.95 – 4.89 (m, 1H), 4.42 (td, *J* = 8.0 Hz, 1.5 Hz, 1H), 1.75 – 1.66 (m, 2H), 1.49 – 1.39 (m, 2H), 1.35 – 1.20 (m, 16H), 0.87 (t, *J* = 4.0 Hz, 3H). **$^{13}$C NMR** (100 MHz, $CDCl_3$): *δ* 136.9, 116.7, 87.7, 83.7, 63.3, 62.6, 37.8, 32.1, 29.8, 29.8, 29.7, 29.7, 29.5, 29.4, 25.3, 22.8, 14.3. **HRMS-DCI** ($CH_4$): *m/z* calcd for $C_{17}H_{31}O_2$ [M+H]$^+$: 267.2324 found: 267.2330. **FTIR** (cm$^{-1}$) (neat): *ν* 3277, 2920, 2852, 1466, 1268, 1146, 1015, 985, 927, 697.

### Supplementary note 5. Spectral data corresponding to the NMR characterization of DACone reaction products with *Nα*-Acetyl Lysine and N-Acetyl Cysteine.

***N$_α$*-Acetyl-L-Lysine:** $^1$H-NMR (700 MHz, $H_2O$): *δ* [ppm] 4.13 Hα (dd, 1H,Hα, $J_{Hα,Hβ}$=8.5 Hz); 2.79 Hε (d, 1H,Hβ, $J_{Hα,Hβ}$=8.5 Hz); 2.02 $H_{CH3}$, (s, 3H, $CH_3$); 1.78 Hβ, (m, 1H, Hβ); 1.67 Hβ, (m, 1H, Hβ); 1.55 $H_δ$ (m, 2H, $H_δ$); 1.37 Hγ (m, 2H, Hγ). $^{13}$C NMR (700 MHz, $H_2O$): *δ* [ppm] 57.94 Cα; 42.83 Cε; 24.70 $C_{CH3}$; 34.05 Cβ; 34.02 Cβ; 31.85 Cδ; 25.11 Cγ.

***N$_α$*-Acetyl L-Lysine modified by DACone 8:** $^1$H-NMR (700 MHz, $H_2O$): *δ* [ppm] 8.03 H (d, 1H,H, *J* = 13.52 Hz); 7.93 H (d, 1H,H, *J*=12.81 Hz); 5.46 H (d, 1H,H, *J*=13.54 Hz); 4.13 Hα (m, 1H, Hα); 3.35 Hε (t, 1H, Hε); 3.22 Hε (t, 1H, Hε); 2.42 H4 (t, 2H, H); 2.00 $H_{CH3}$ (s, 1H, $CH_3$); 1.80 Hβ (m, 1H, Hβ); 1.66 Hβ (m, 1H, Hβ); 1.61 Hδ (m, 2H, Hδ); 1.56 H3 (m, 2H, H3); 1.42 H2 (m, 2H, H2); 1.38 Hγ (m, 2H, Hγ); 0.90 H (t, 3H, H1). $^{13}$C NMR (700MHz, $H_2O$): *δ* [ppm] 162.79 $C_9$; 166.44 $C_9$; 102.38 $C_8$; 57.71 Cα; 51.45 Cε; 45.95 Cε; 32.08 $C_4$; 24.5 $C_{CH3}$; 33.93 Cβ; 33.90 Cβ; 29.70 Cδ; 32.08 $C_3$; 24.28 $C_2$; 25.04 Cγ; 15.42 $C_1$

***N*-Acetyl-L-Cysteine:** $^1$H-NMR (700 MHz, $H_2O$): *δ* [ppm] 4.37 Hα (t, 1H, Hα); 2.91 Hβ (d, 1H, Hβ); 2.90 Hβ (d, 1H, Hβ); 2.05 $H_{CH3}$, (s, 3H, $CH_3$). $^{13}$C NMR (700MHz, $H_2O$): δ [ppm] 57.94 Cα; 42.83 Cε; 24.70 $C_{CH3}$; 34.05 Cβ; 34.02 Cβ; 31.85 C δ; 25.11 Cγ.

***N*-Acetyl-L-Cysteine modified by DACone 8*:*** $^1$H-NMR (700 MHz, $H_2O$): *δ* [ppm] 8.23 H8 (d, 1H (0.25), H8, $J_{H,H}$=15.20 Hz); 7.59 H8 (d, 1H (0.75), H8, $J_{H,H}$=9.98 Hz); 6.51 H9 (d, 1H (0.75), H9, $J_{H,H}$=9.98 Hz); 6.34 H9 (d, 1H (0.25), H9, $J_{H,H}$=15.20 Hz); 4.53 Hα (dd, 1H (0.25), Hα); 4.46 Hα (dd, 1H (0.75),Hα); 3.45 Hβ (dd, 1H (0.25), Hβ); 3.37 Hβ (dd, 1H (0.75), Hβ); 3.29 Hβ (dd, 1H (0.25), Hβ); 3.19 Hβ (dd, 1H (0.75), Hβ); 2.49 H4 (t, 2H (0.25), H4); 2.45 H4 (t, 2H (0.75), H4); 2.02 $H_{CH3}$, (s, 3H, $CH_3$); 1.59 H3 (q, 2H (0.25), H3); 1.56 H3 (q, 2H (0.75), H3); 1.43 H2 (sx, 2H (0.25), H2); 1.42 H2 (sx, 2H (0.75), H2); 0.90 H1 (t, 3H (0.25), H1); 0.89 H1 (t, 3H (0.75), H1). $^{13}$C NMR (700 MHz, $H_2O$): *δ* [ppm] 161.24 C1; 157.68 C8 ; 125.06 C9; 127.65 C9; 56.79 Cα; 57.57 Cα; 38.00 Cβ; 41.81 Cβ; 38.02 Cβ; 41.79 Cβ; 20.74 C4; 20.77 C4; 24.61 $C_{CH3}$; 31.78 C3; 24.12 C2; 15.36 C1.

### Supplementary note 6. NMR spectra of novel compounds

$^1$H NMR spectrum ($CD_3CN$, 300 MHz) of compound **7**

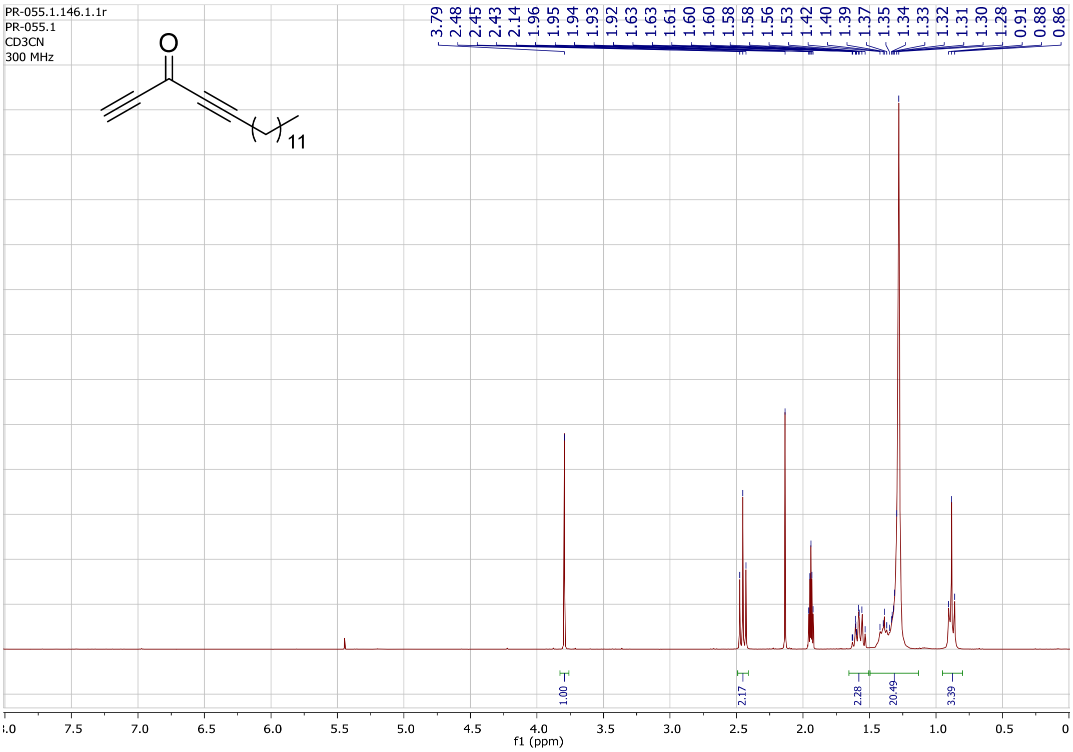

$^{13}$C NMR spectrum ($CD_3CN$, 75 MHz) of compound **7**

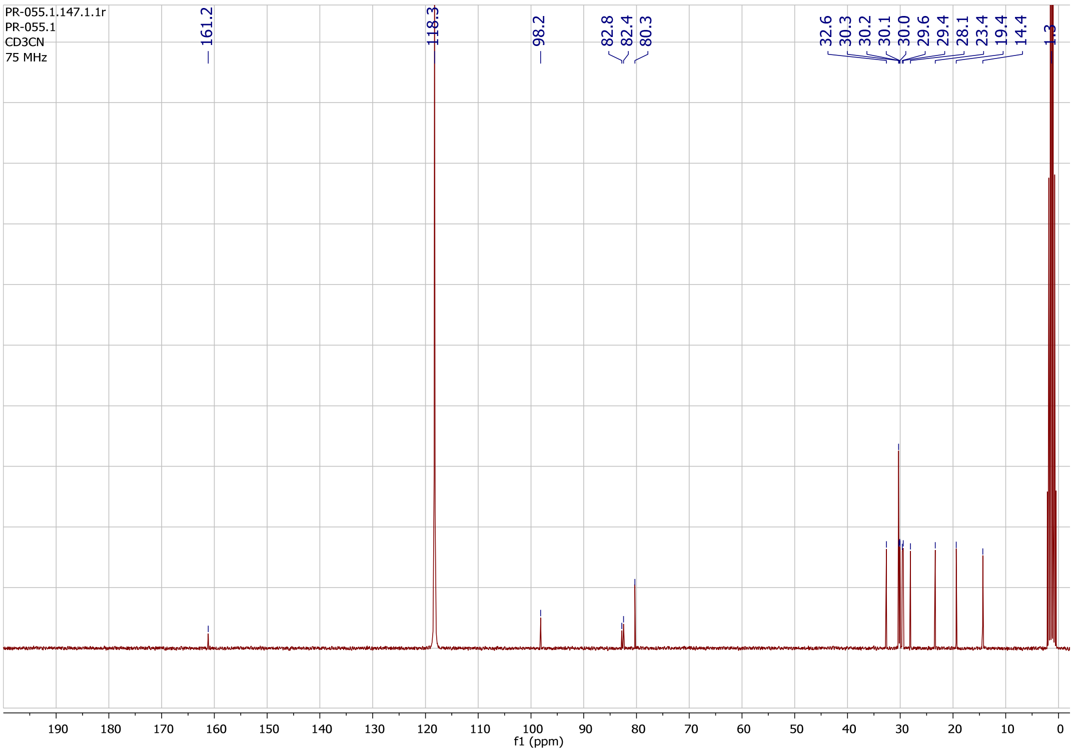

$^1$H NMR spectrum ($CDCl_3$, 300 MHz) of compound **CBD513**

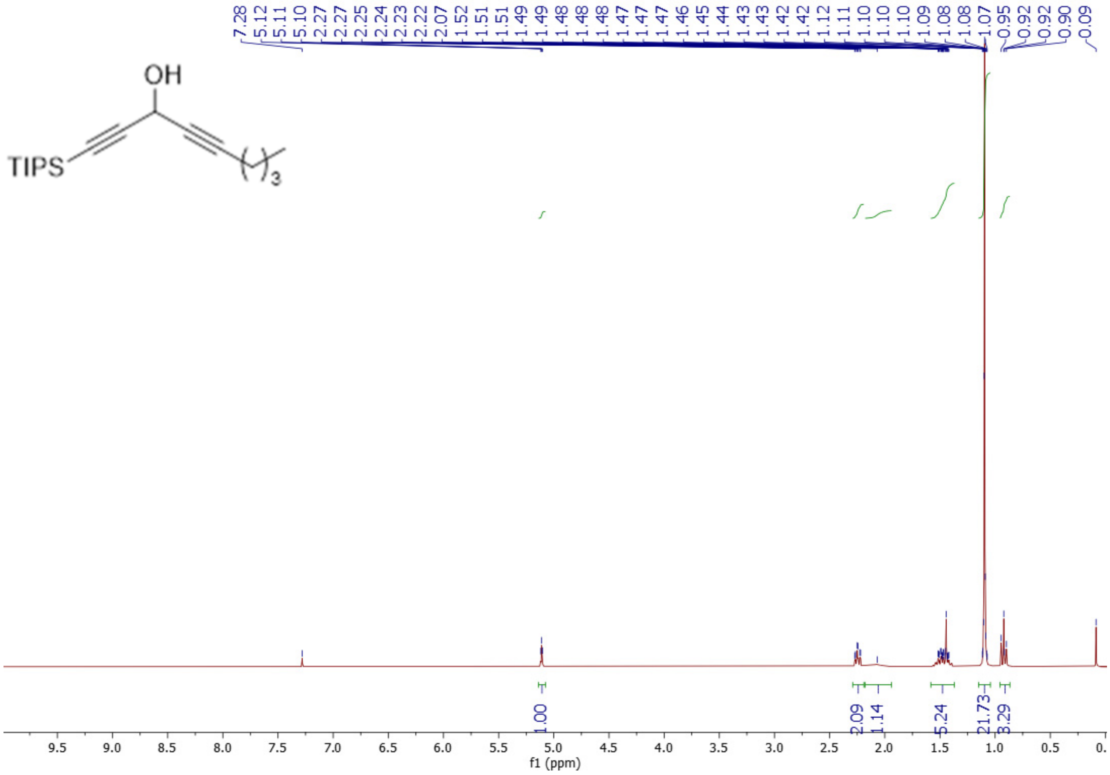

$^{13}$C NMR spectrum ($CDCl_3$, 75 MHz) of compound **CBD513**

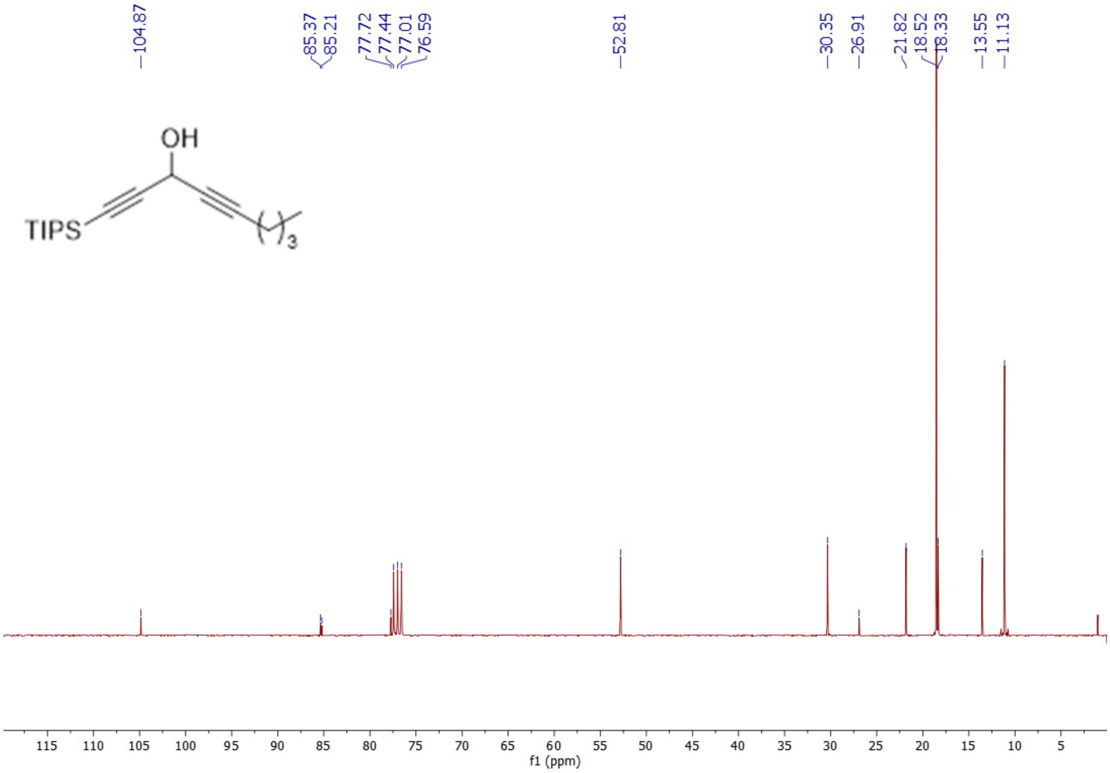

$^1$H NMR spectrum ($CDCl_3$, 300 MHz) of compound **CBD515**

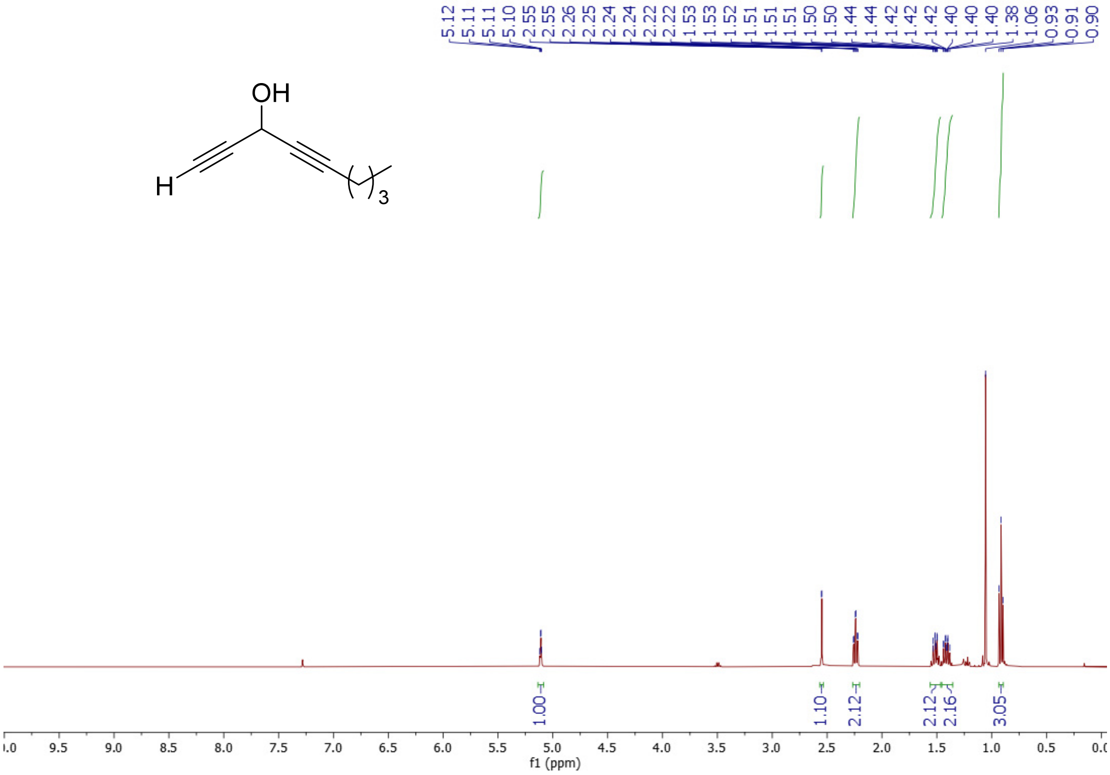

$^{13}$C NMR spectrum ($CDCl_3$, 101 MHz) of compound **CBD515**

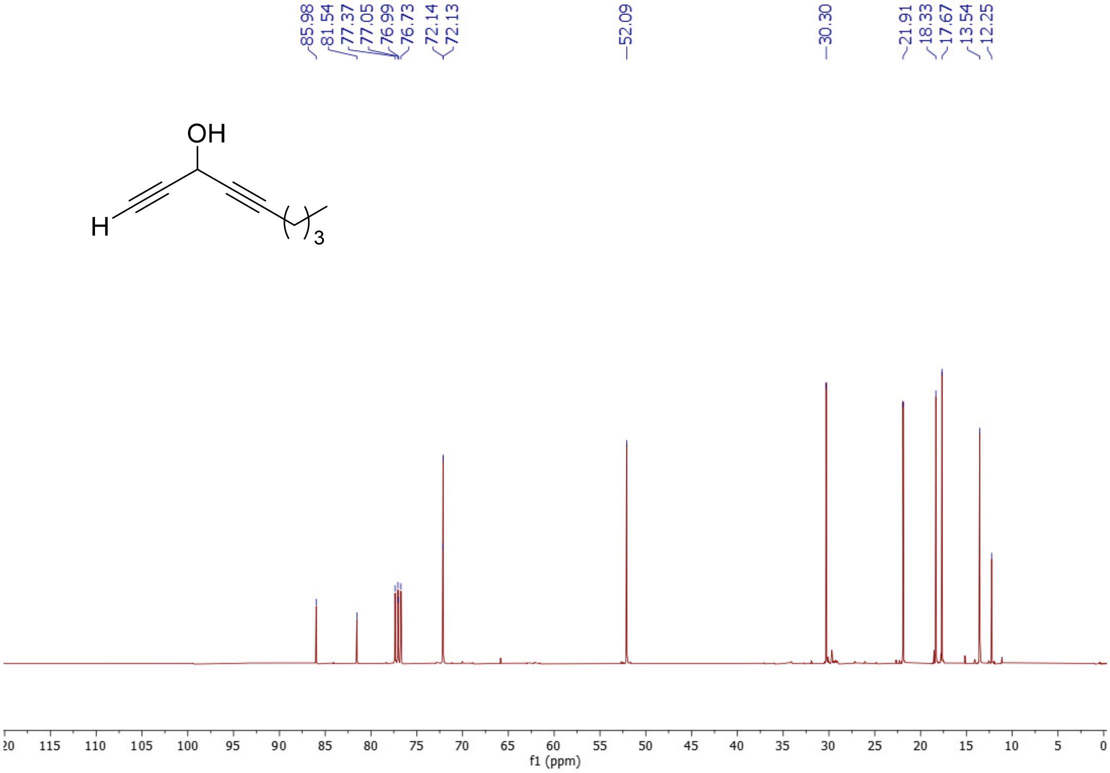

$^{1}H$ NMR spectrum ($CDCl_3$, 300 MHz) of compound **8**

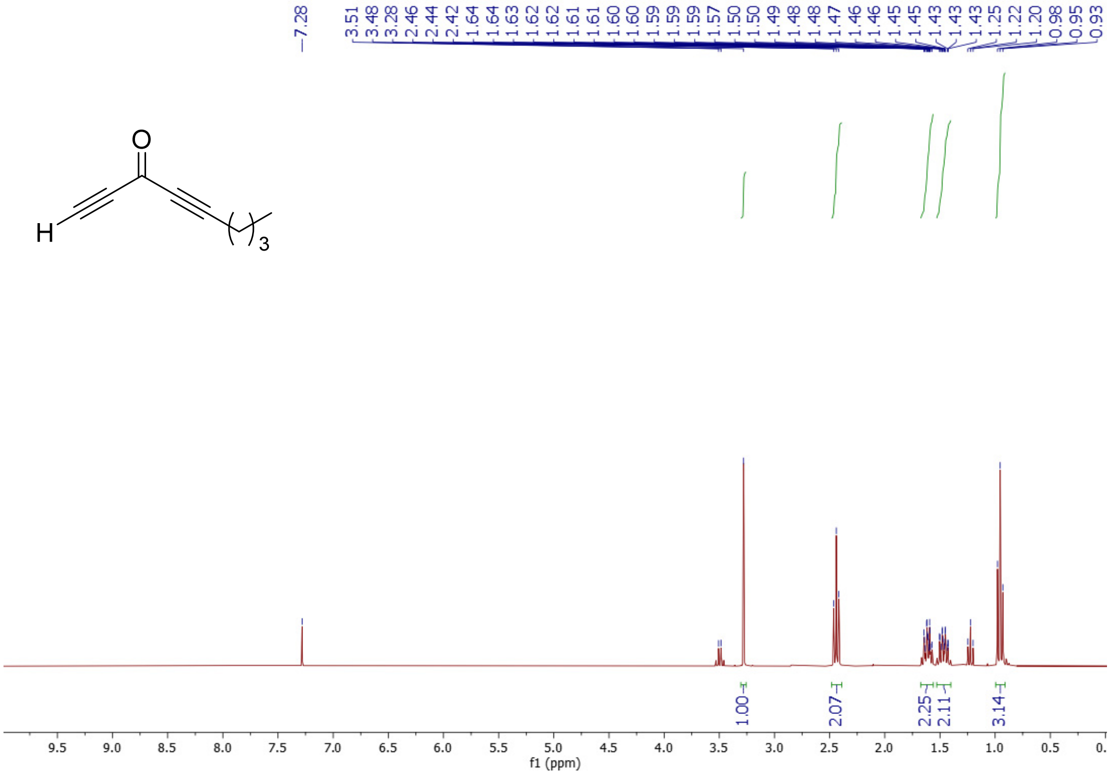

$^{13}C$ NMR spectrum ($CDCl_3$, 75 MHz) of compound **8**

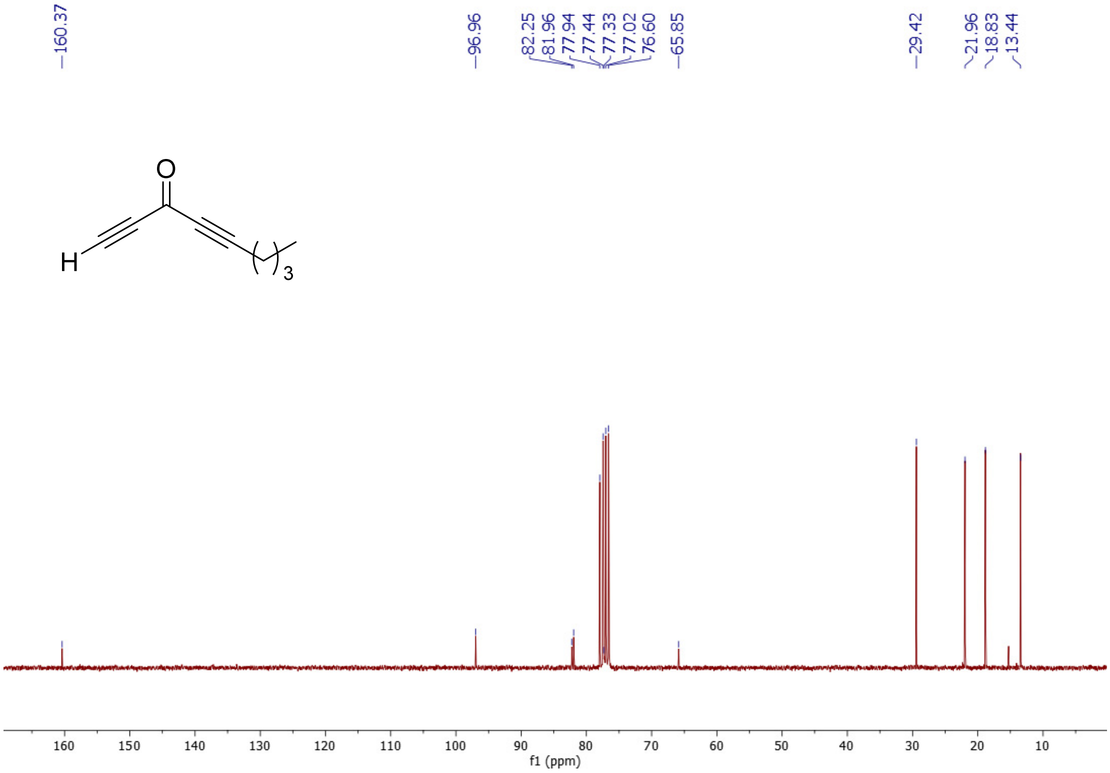

$^{1}H$ NMR spectrum ($CD_3CN$, 300 MHz) of compound **10**

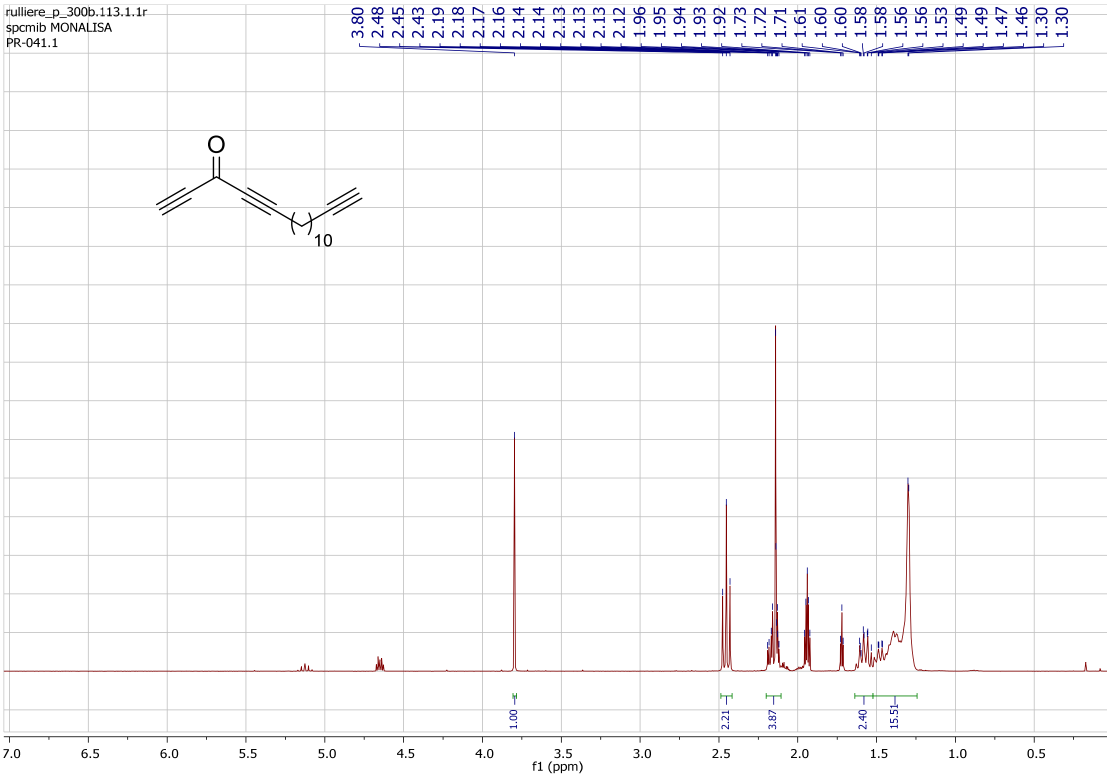

$^{13}C$ NMR spectrum ($CD_3CN$, 75 MHz) of compound **10**

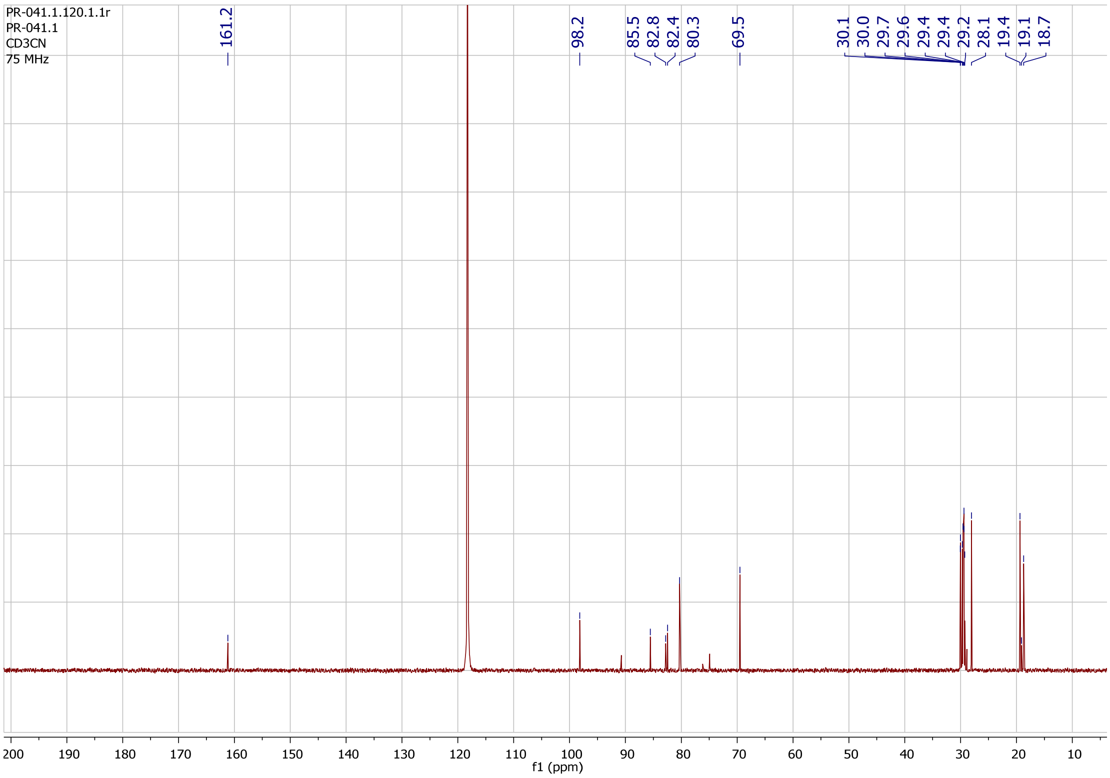

$^1$H NMR spectrum ($CDCl_3$, 400 MHz) of compound **CBD514**

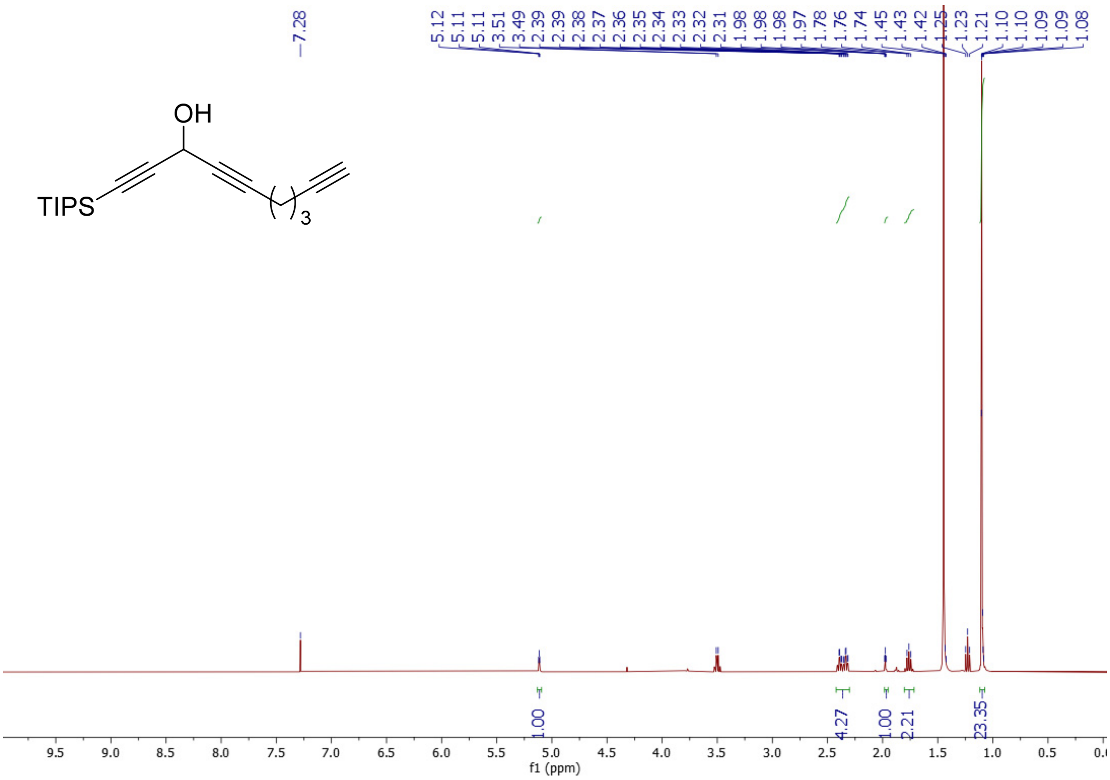

$^{13}$C NMR spectrum ($CDCl_3$, 75 MHz) of compound **CBD514**

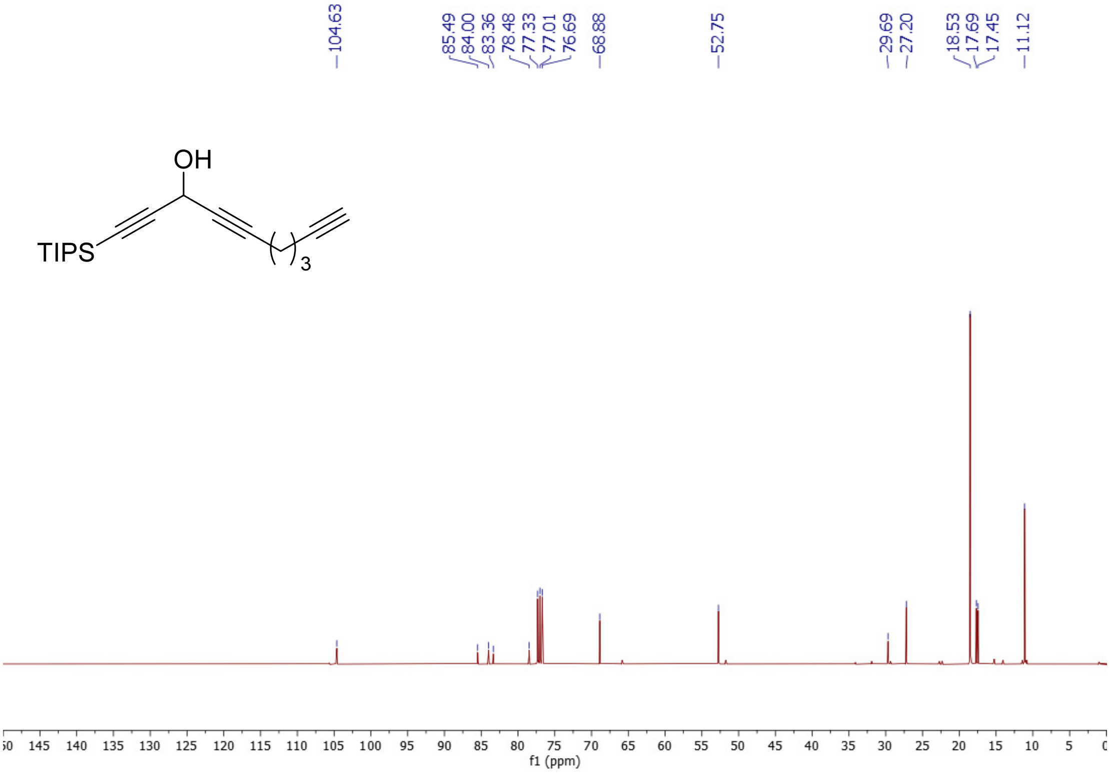

$^1H$ NMR spectrum ($CDCl_3$, 300 MHz) of compound **CBD516**

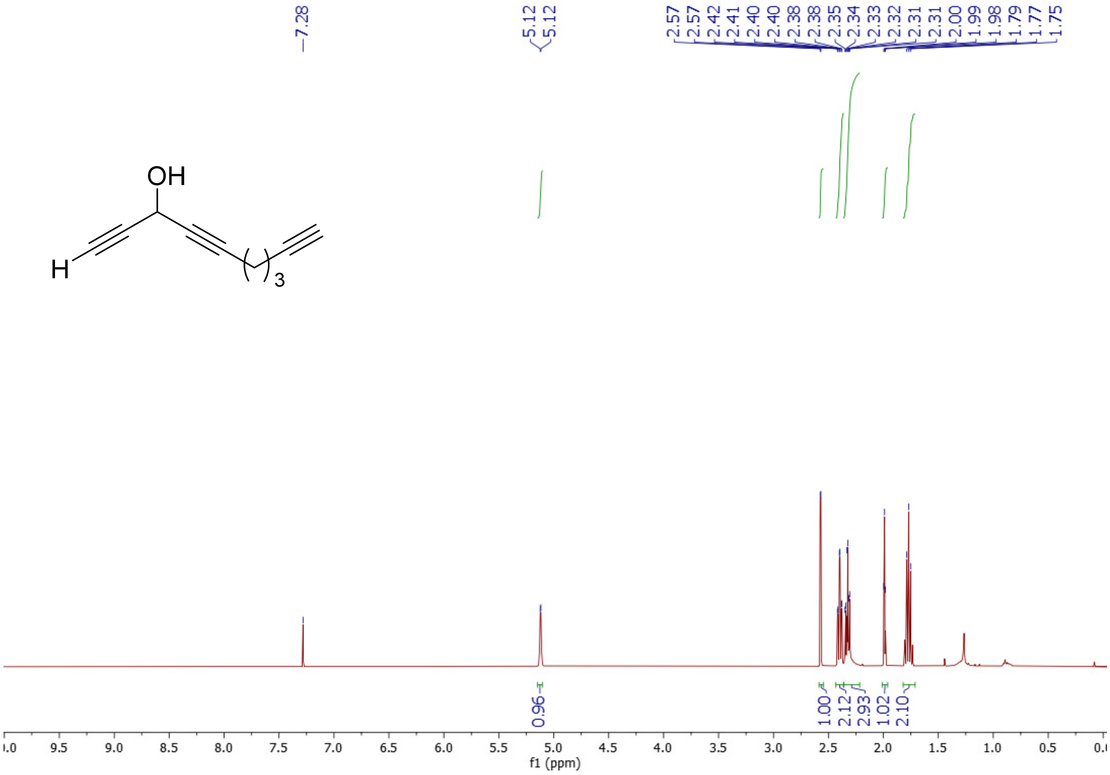

$^{13}C$ NMR spectrum ($CDCl_3$, 75 MHz) of compound **CBD516**

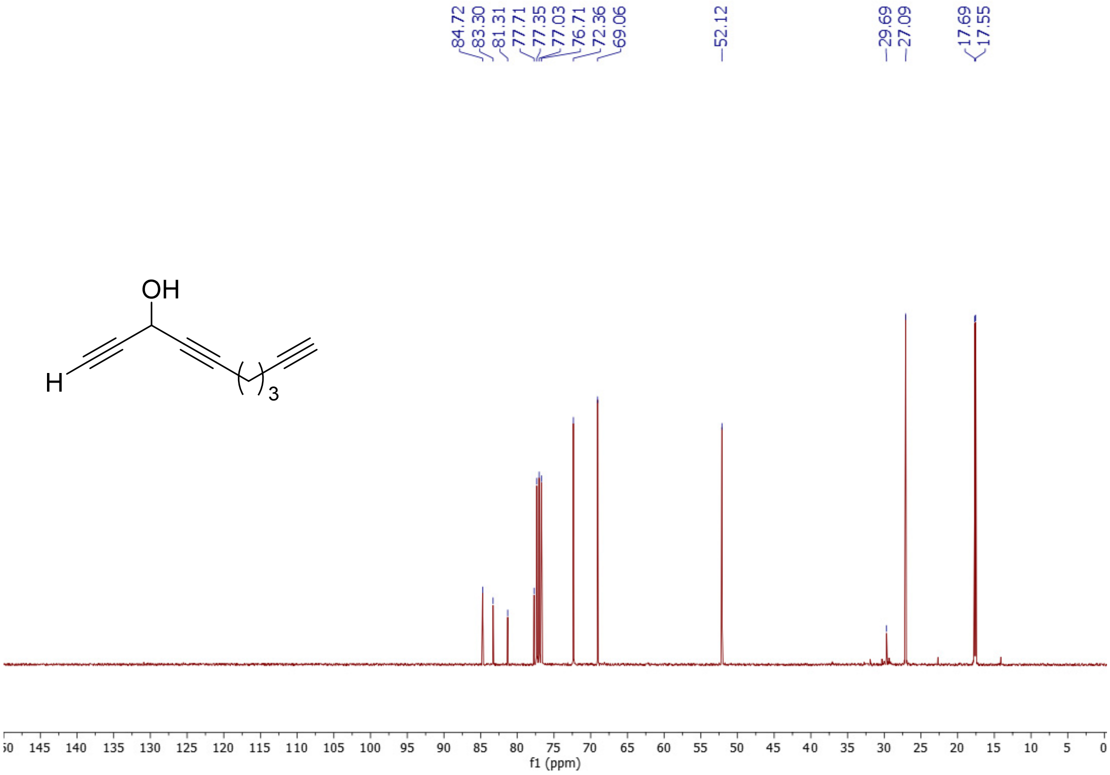

[1]H NMR spectrum ($CDCl_3$, 300 MHz) of compound **11**

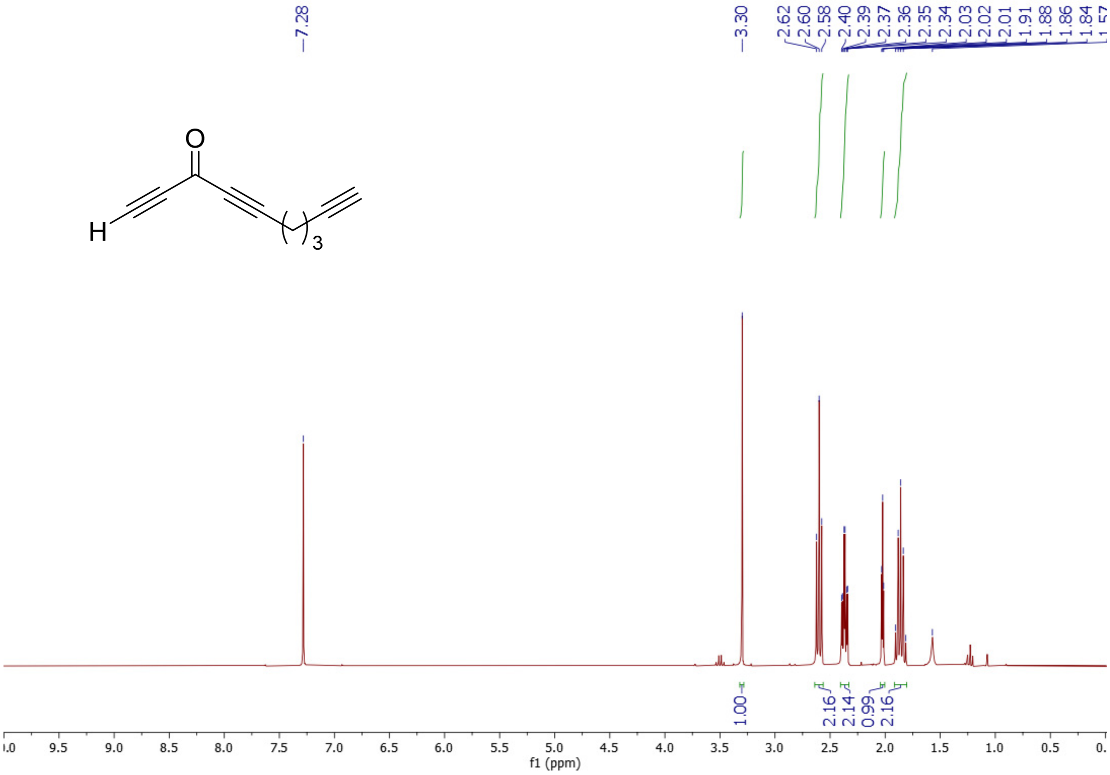

[13]C NMR spectrum ($CDCl_3$, 75 MHz) of compound **11**

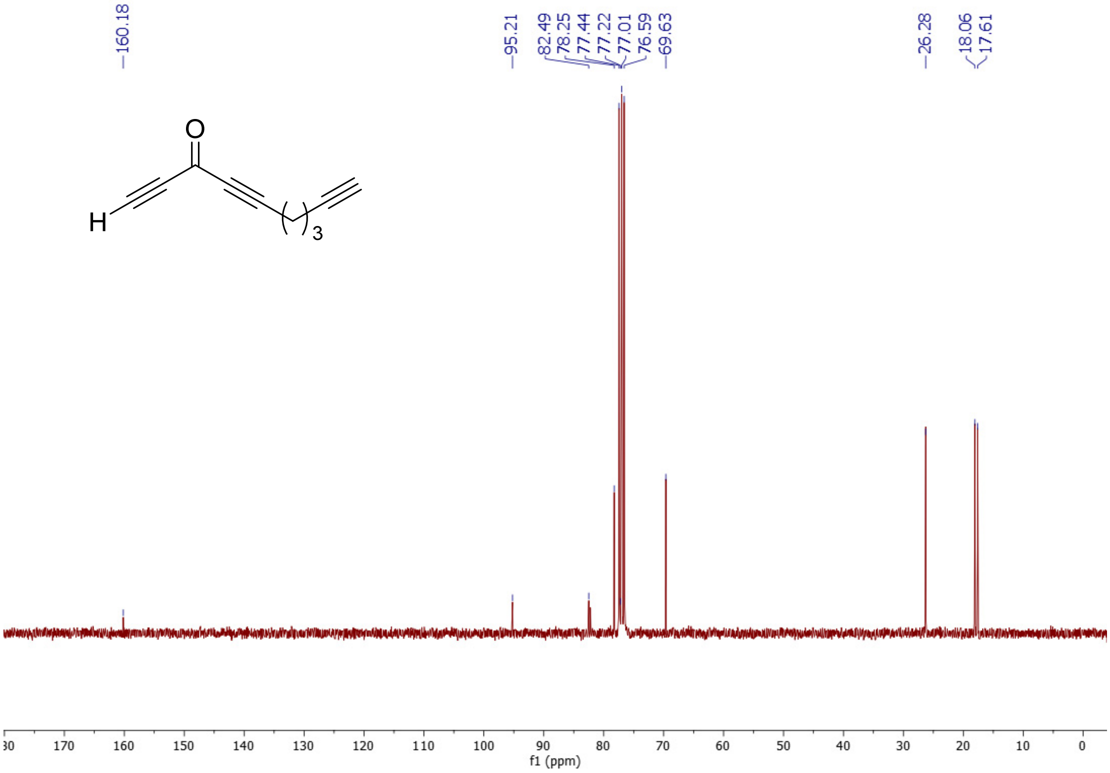

### $^{1}H$ NMR spectrum ($CDCl_3$, 300 MHz) of compound **MVB39**

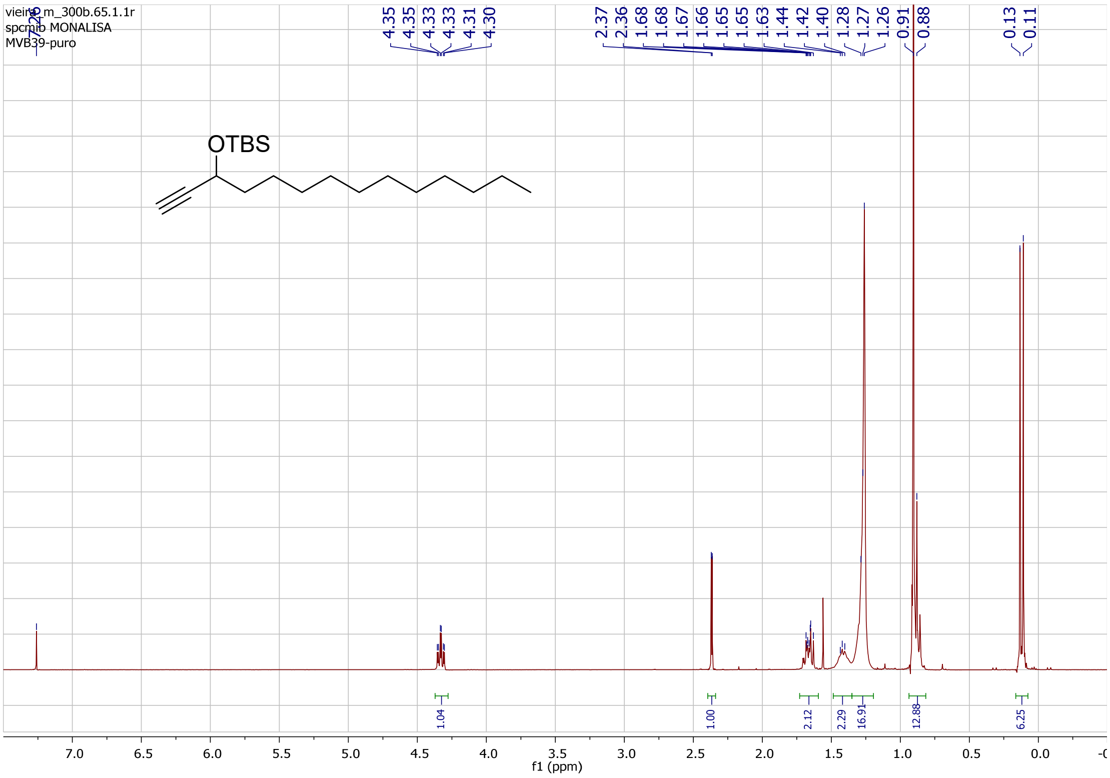

### $^{13}C$ NMR spectrum ($CDCl_3$, 75 MHz) of compound **MVB39**

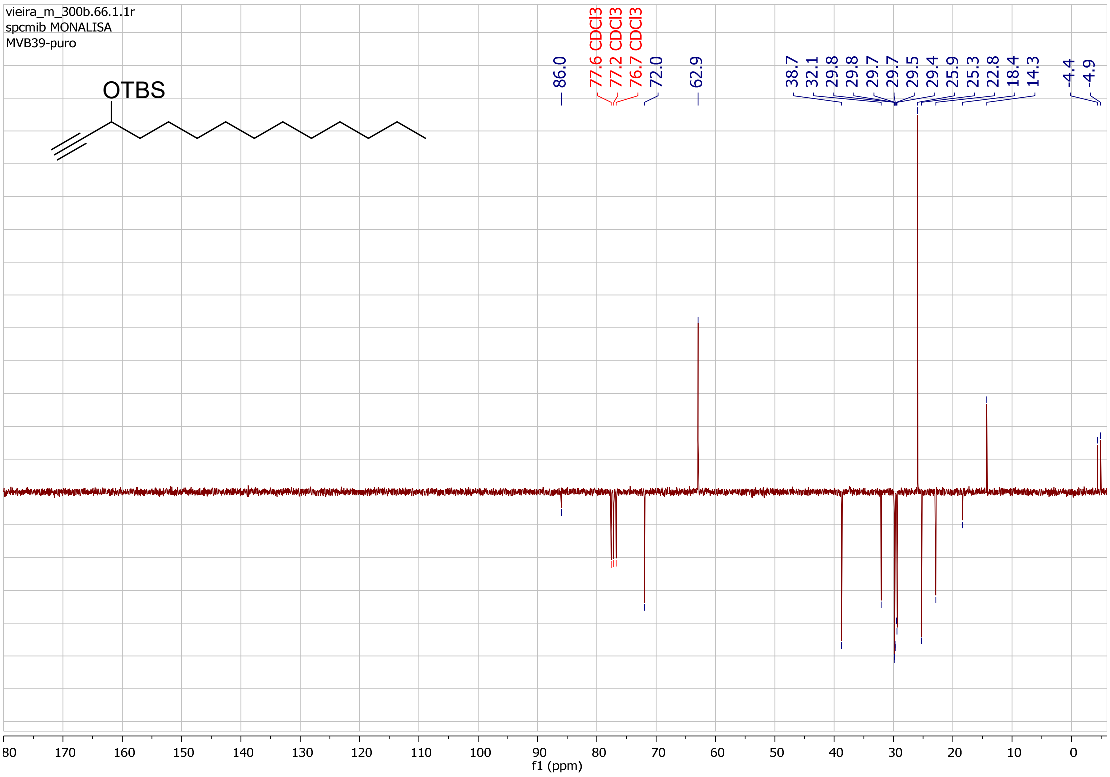

$^{1}H$ NMR spectrum ($CDCl_3$, 400 MHz) of compound **MVBL05**

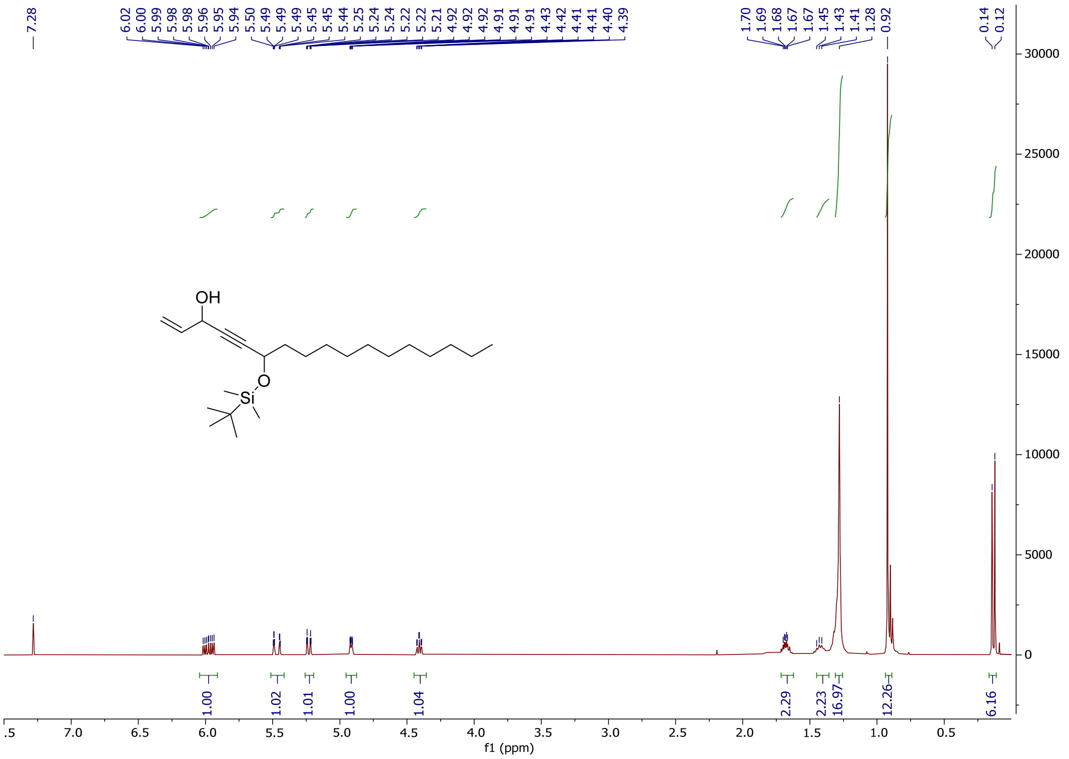

$^{13}C$ NMR spectrum ($CDCl_3$, 100 MHz) of compound **MVBL05**

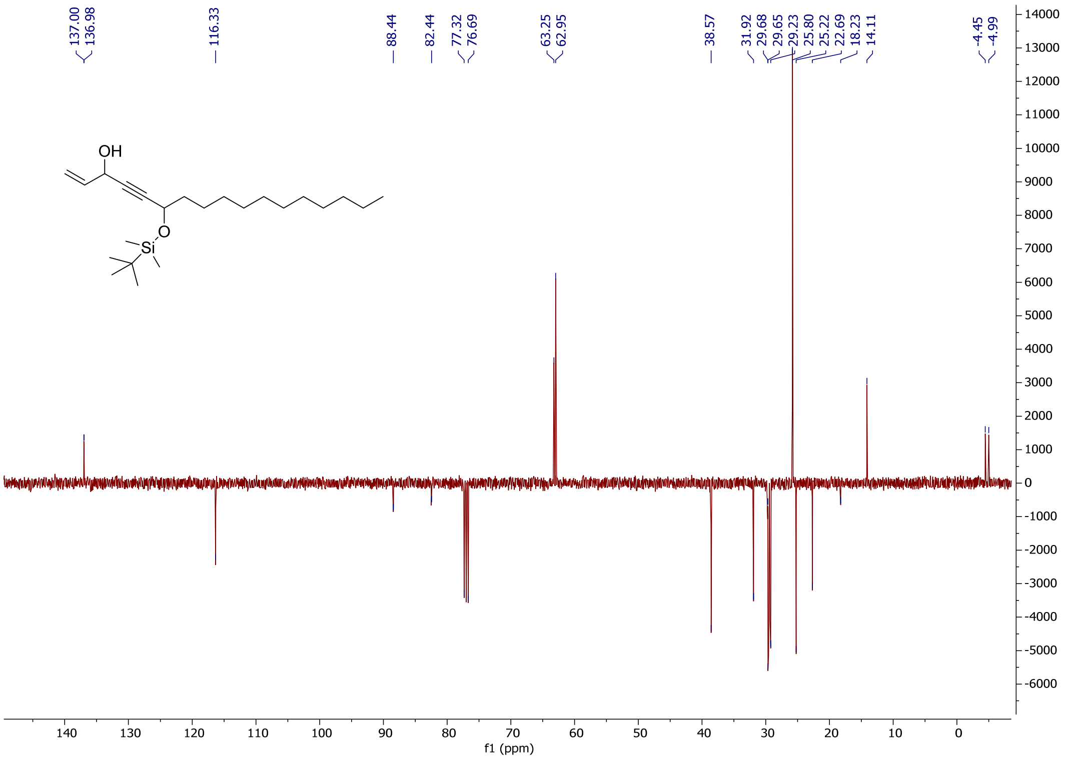

[1]H NMR spectrum ($CDCl_3$, 400 MHz) of compound **12**

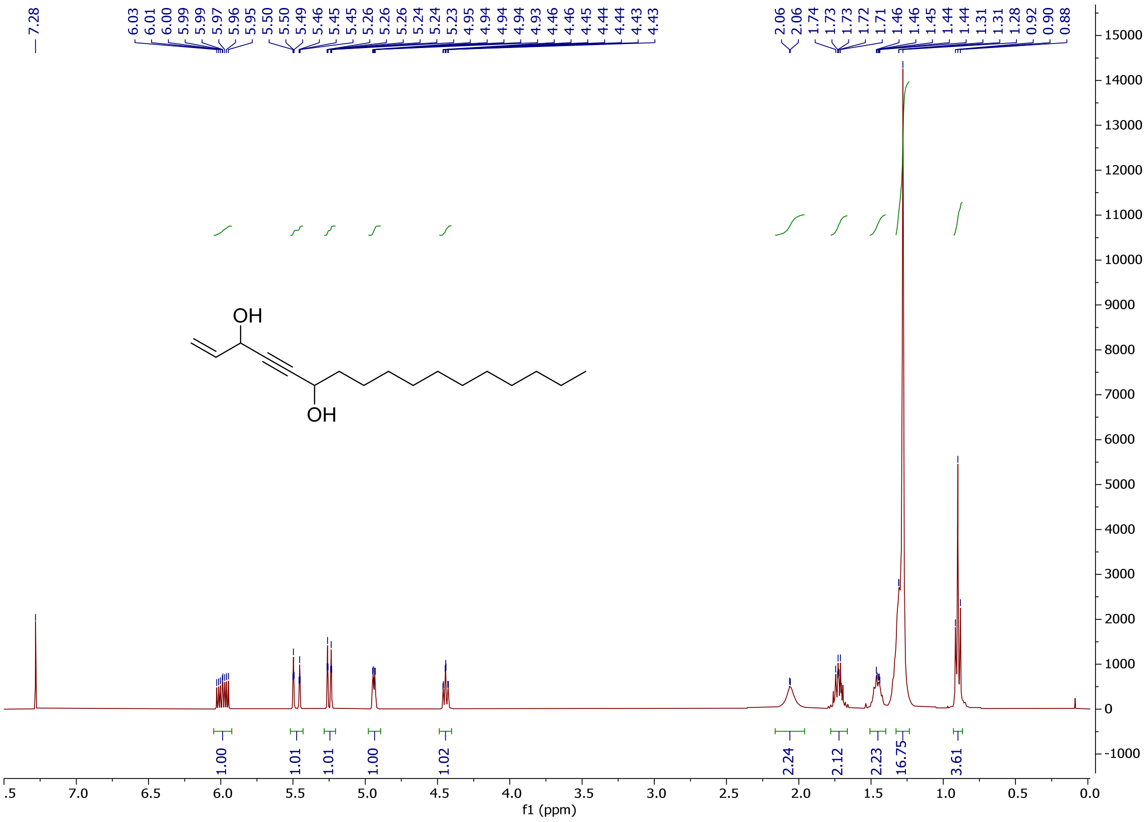

[13]C NMR spectrum ($CDCl_3$, 100 MHz) of compound **12**

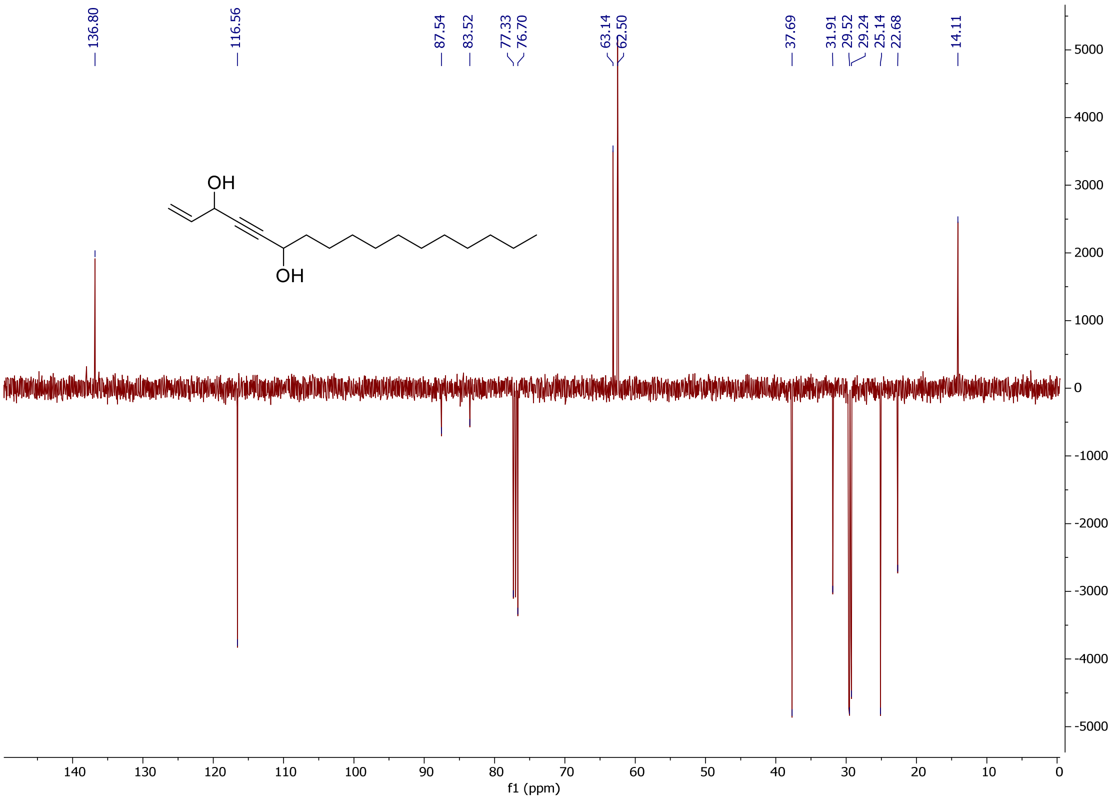
